# Transcriptomic signatures associated with regional cortical thickness changes in Parkinson’s disease

**DOI:** 10.1101/2020.06.19.158808

**Authors:** Arlin Keo, Oleh Dzyubachyk, Jeroen van der Grond, Jacobus J. van Hilten, Marcel J. T. Reinders, Ahmed Mahfouz

**Affiliations:** Leiden Computational Biology Center, Leiden University Medical Center, Leiden, the Netherlands; Delft Bioinformatics Lab, Delft University of Technology, Delft, the Netherlands; Department of Radiology, Leiden University Medical Center, Leiden, the Netherlands; Department of Neurology, Leiden University Medical Center, Leiden, the Netherlands; Department of Human Genetics, Leiden University Medical Center, Leiden, the Netherlands

**Keywords:** cortical thickness, neurodegenerative diseases, neuroimaging data, imaging-genetics, gene expression analysis

## Abstract

Cortical atrophy is a common manifestation in Parkinson’s disease, particularly in later disease stages. Here, we investigated patterns of cortical thickness using T1-weighted anatomical MRI data of 149 Parkinson’s disease patients and 369 controls. To elucidate the molecular underpinnings of cortical thickness changes in Parkinson’s disease, we performed an integrated analysis of brain-wide healthy transcriptomic data from the Allen Human Brain Atlas and neuroimaging features. For this purpose, we used partial least squares regression to identify gene expression patterns correlated with cortical thickness changes. In addition, we identified gene expression patterns underlying the relationship between cortical thickness and clinical domains of Parkinson’s disease. Our results show that genes whose expression in the healthy brain is associated with cortical thickness changes in Parkinson’s disease are enriched in biological pathways related to sumoylation, regulation of mitotic cell cycle, mitochondrial translation, DNA damage responses, and ER-Golgi traffic. The associated pathways were highly related to each other and all belong to cellular maintenance mechanisms. The expression of genes within most pathways was negatively correlated with cortical thickness changes, showing higher expression in regions associated with decreased cortical thickness (atrophy). On the other hand, sumoylation pathways were positively correlated with cortical thickness changes, showing higher expression in regions with increased cortical thickness (hypertrophy). Our findings suggest that alterations in the balanced interplay of these mechanisms play a role in changes of cortical thickness in Parkinson’s disease and possibly influence motor and cognitive functions.

## Introduction

Parkinson’s disease is a neurodegenerative disorder characterized by a progressive loss of dopaminergic and non-dopaminergic neurons in the brain and peripheral and autonomic nervous system (Hirsch *et al*., 2012). Cortical atrophy occurs during the later disease stages and has been associated with cognitive decline, including executive, attentional, memory, and visuospatial deficits (Aarsland *et al*., 2017; Wilson *et al*., 2019). Although MRI studies of patient brains have tried to link regional cortical atrophy to clinical features of the disease (Rosenberg-Katz *et al*., 2016; Wang *et al*., 2016; Chen *et al*., 2017; Li *et al*., 2018; Zheng *et al*., 2019), little is known about the pathobiology that underlies the selective cortical vulnerability in Parkinson’s disease.

Analyzing the transcriptome in vulnerable cortical regions may aid in better understanding the underlying molecular mechanisms of atrophy in Parkinson’s disease. Although gene expression data of human post-mortem Parkinson’s disease brains is available, most findings relate to studies that focused only on one or few coarse brain regions (Oerton and Bender, 2017). To perform whole brain analysis of both gene expression and imaging data, studies turn to the Allen Human Brain Atlas (AHBA), a high resolution gene expression atlas covering the entire brain of six adult donors without any history of neurological disorders (Hawrylycz *et al*., 2015; Arnatkevičiūtė *et al*., 2019). The AHBA has been combined with functional MRI data of Parkinson’s disease patients and revealed that the regional expression of *MAPT*, but not *SNCA*, correlated with the loss of regional connectivity (Rittman *et al*., 2016). Using a similar approach, correlations were identified between a cortical atrophy pattern and the regional expression of 17 genes implicated in Parkinson’s disease (Freeze *et al*., 2018). Although both studies used spatial transcriptomics to explore gene expression across the whole brain, they only analyzed the expression of a limited set of genes that are of interest to Parkinson’s disease, e.g. those that are known as genetic risk factors.

To investigate the relationship between high dimensional genome-wide expression patterns and imaging data, multivariate analysis methods are required. Partial least squares (PLS) regression has been used to perform simultaneous analysis of brain-wide gene expression from the AHBA and neuroimaging data of adolescents, healthy adults, and Huntington’s disease (Vértes *et al*., 2016; Whitaker *et al*., 2016; McColgan *et al*., 2017). The PLS approach allows the linking of multiple predictor variables (genes) and multiple response variables (imaging features) and deals with multicollinearity by projecting variables to a smaller set of components that are maximally correlated between both datasets. Thus, PLS is an attractive model to identify gene expression patterns associated with imaging features.

Here, we exploited PLS regression to find transcriptomic signatures that are related to changes in cortical thickness (CT) in Parkinson’s disease. MRI data was obtained from patients and age-matched controls to find CT changes across all cortical regions. Gene expression samples from healthy donors in the AHBA were anatomically mapped to the cortical regions to find brain-wide gene expression patterns predictive of the CT changes observed in Parkinson’s disease patients. In addition, we assessed the relationships between CT and clinical scores in Parkinson’s disease patients and used a second PLS model to find expression patterns associated with these relationships across all cortical regions. With these models we address three research questions: 1) Which cortical regions show CT changes in Parkinson’s disease, 2) Which genes and biological pathways show expression patterns associated with these regional changes, and 3) Which molecular mechanisms underlie the relationships between CT and clinical scores in Parkinson’s disease. To answer these questions, we explored the whole transcriptome in cortical regions of the healthy brain to find expression signatures predictive of imaging features in Parkinson’s disease.

## Material and methods

### MRI data acquisition

MRI images of 149 Parkinson’s disease patients (mean age = 64.8 years; 65.7% male) were obtained from a cross-sectional cohort study and is part of the ‘PROfiling PARKinson’s disease’ (PROPARK) study (de Schipper *et al*., 2017). Parkinson’s disease patients were recruited from the outpatient clinic for Movement Disorders of the Department of Neurology of the Leiden University Medical Center and nearby university and regional hospitals. All participants fulfilled the United Kingdom Parkinson’s Disease Society Brain Bank criteria for idiopathic Parkinson’s disease (Gibb and Lees, 1988); written consent was obtained from all participants. Three-dimensional T1-weighted anatomical images were acquired on a 3 Tesla MRI scanner (Philips Achieva, Best, the Netherlands) using a standard 32-channel whole-head coil. Acquisition parameters were: repetition time = 9.8 ms, echo time = 4.6 ms, flip angle = 8°, field of view 220 × 174 × 156 mm, 130 slices with a slice thickness of 1.2 mm with no gap between slices, resulting in a voxel size of 1.15 mm × 1.15 mm × 1.20 mm.

Three-dimensional T1-weighted images from 369 controls (mean age = 65.7 years; 48.1% male) were acquired in a different cohort (Altmann-schneider *et al*., 2012), where all imaging was performed on a whole body 3 Tesla MRI scanner (Philips Medical Systems, Best, the Netherlands), using the following imaging parameters: TR = 9.7 ms, TE = 4.6 ms, FA = 8°, FOV = 224 × 177 × 168 mm. The anatomical images covered the entire brain with no gap between slices resulting in a nominal voxel size of 1.17 × 1.17 × 1.4 mm. Acquisition time was approximately 5 min.

### Cortical thickness changes in segmented cortical regions

CT in cortical regions of Parkinson’s disease patients and controls was determined using cortical parcellation implemented in FreeSurfer version 5.3.0 (Fischl and Dale, 2000). The FreeSurfer algorithm automatically parcellates the cortex and assigns a neuroanatomical label to each location on a cortical surface model based on probabilistic information. The parcellation scheme of the Desikan–Killiany atlas was used to divide the cortex into 34 regions per hemisphere (Desikan *et al*., 2006).

To assess CT changes between patients (149) and controls (369), a two-tailed *t*-test assuming unequal variances was applied in SPSS Statistics version 23. *P*-values were corrected for multiple testing across 68 cortical regions using the Benjamini-Hochberg (BH) method. A two-tailed *t*-test was also used to assess CT differences between the left and right hemisphere for each one of the 34 cortical regions, with *P*-values being BH-corrected across the 34 cortical regions.

### Clinical scores

All patients underwent standardized assessments, and an evaluation of demographic and clinical characteristics (de Schipper et al., 2017). MDS-UPDRS is a clinical rating scale consisting of four parts: I) Non-motor Experiences of Daily Living; II) Motor Experiences of Daily Living; III) Motor Examination; IV) Motor Complications (Goetz et al., 2008). UPDRSTOTSCR is the total score of all four parts. The SENS-PD scale is a composite score of non-dopaminergic symptoms (van der Heeden et al., 2016), LED is the levodopa equivalent dose (Tomlinson et al., 2010), and MMSE is the mini-mental state examination (Folstein et al., 1975).

### Relationship between CT and clinical scores

We used CT data and clinical scores from 149 Parkinson’s disease patients to determine the relationships between CT and clinical domains. We selected 9 clinical features with numeric (non-nominal) values for which scores were available for 82-123 patients: AGEONSET, SENSPDSC, MDS_UPDRS_3, MMSE, LED, MDS_UPDRS_1, MDS_UPDRS_2, MDS_UPDRS_4, and UPDRSTOTSCR (Supplementary Fig. 1).

The correlation between CT and the scores of each clinical feature individually was determined across patients by applying linear regression. To obtain maximum correlation, separate linear regression models were used for each combination of a region and clinical feature:

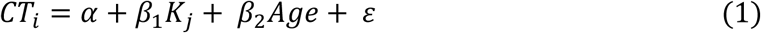

where *CT*_*i*_ is the CT of one region *i* across patients, *K*_*j*_ is the score of one clinical feature *j* across patients. *Age* is taken into account to correct for the age of patients. *α* is the background term, *β*_1_ is the regression coefficient for *K*_*j*_, *β*_2_ is the regression coefficients for *Age*, and *ε* is the residual. The regression coefficient *β*_1_ was used to determine the relationship between CT and clinical domain scores, and assessed for statistical significance where *P*-values were BH-corrected for 34 regions and 9 clinical features (*t*-test, H_0_: *β*_1_ = 0, *P* < 0.05).

### Mapping transcriptomic data to cortical regions

We downloaded normalized gene expression data from the Allen Human Brain Atlas (AHBA; http://human.brain-map.org/), a microarray data set of 3,702 anatomical brain regions from six non-neurological individuals (5 males and 1 female, mean age 42, range 24–57 years (Hawrylycz *et al*., 2015)). To analyze the transcriptome in the cortical regions, we used the mapping of AHBA samples to cortical regions in neuroimaging data proposed in (Arnatkevičiūtė *et al*., 2019), where they applied Freesurfer on T1 MRIs of the six donors in the AHBA to segment the cortical regions according to the Desikan-Killiany atlas. AHBA samples were mapped to 34 cortical regions from the left hemisphere, since for only two out of six brains samples were collected from both hemispheres and for four brains they only sampled from the left hemisphere. By only analyzing the left hemisphere, we assumed that there are small to no differences in gene expression between the left and right hemisphere (Hawrylycz *et al*., 2015). Samples were assigned to a segmented cortical region when their MNI coordinates corresponds to a voxel within a parcel, including samples that are up to 2 mm away from any voxel in the parcel. In total 1,284 samples from the AHBA were assigned to the 34 cortical regions.

### Partial least squares (PLS) model-1 and model-2

We used PLS regression (R-package *pls* 2.7) to find gene expression patterns across the 34 cortical regions that are predictive of gray matter atrophy and possibly their relationship to scores of nine clinical domains (Supplementary Methods). PLS regression and principal component analysis regression are both methods where the original measurements are projected to latent variables to study the data in reduced dimensions (Fig. 1A). PLS however, projects variables from each dataset to latent variables such that they are maximally correlated between two datasets *X* and *Y* (Fig. 1B). In this study, the predictor *X* is a gene expression matrix of 34 regions (*n*) in the left hemisphere and all 20,017 genes (*m*) and is used to predict imaging variables (*p*) in the same set of 34 cortical regions. For each cortical region and each gene, expression levels were averaged across samples that fall within that cortical region and then averaged across the six donors from the AHBA, such that the input matrix of predictor variables contains one expression value for every gene per cortical region. We implemented two PLS models (Fig. 1C): one single-response PLS model, *model-1*, to predict CT changes, measured as the *t*-statistics of ΔCT between Parkinson’s disease patients and controls, and one multi-response PLS model, *model-2*, to predict the correlation between CT and clinical scores in Parkinson’s disease patients, measured as the *t*-statistics of the coefficients *β*_1_ in Equation 1.

**Figure 1.**
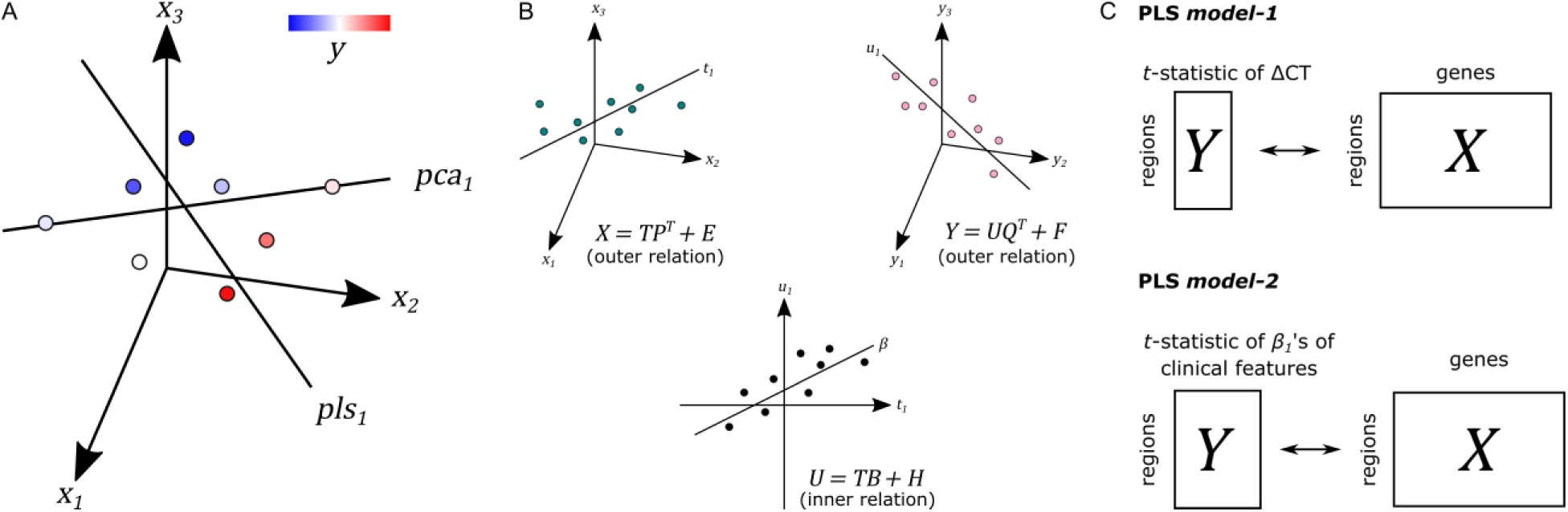
Principal of partial least squares regression (PLS). (A) Principal component analysis (PCA) and PLS project measurements to a new latent space. Unlike PCA, PLS tries to find a latent space that is maximally correlated with another measurement ***y*** from dataset ***Y*** on the same samples. (B) The first latent component ***t***_**1**_ of dataset ***X*** is maximally correlated with the first latent factor ***u***_**1**_ of dataset ***Y. T*** and ***U*** scores determine the outer relations of individual datasets in the model. The coefficient ***β*** determines the inner relation between both datasets X and Y in the model (more details in Supplementary Methods). (C) In PLS *model-1*, we used regional gene expression as input to predict the regional *t*-statistics of ΔCT. Given the PLS model, ***R*** in Equation 5 and 6 in Supplementary Methods is used as gene weights. In PLS *model-2*, we used the same input to predict the *t*-statistics of correlation coefficients ***β***_**1**_ of clinical features from Equation 1.

### Pathway enrichment

Pathway enrichment analysis was done using gene set enrichment analysis (GSEA) and 2,225 pathways from the Reactome database in ReactomePA R-package version 1.28. Genes were ranked based on their weights to each PLS component; *R* in Equation 5 and 6 in Supplementary Methods. Pathways were significant when the FDR-adjusted *P* < 0.05.

### Data and code availability

Transcriptomic data from the AHBA is available at http://human.brain-map.org/. All scripts were run in R version 1.6 and can be found online at https://github.com/arlinkeo/pd_pls.

## Results

### CT changes between Parkinson’s disease patients and controls

We analyzed CT changes between Parkinson’s disease patients and healthy controls (ΔCT) as a measure for gray matter loss (Fig. 1C). Each of the 68 cortical regions from both hemispheres was assessed, for which ΔCT was statistically significant in 10 cortical regions (*t*-test, BH-corrected *P* < 0.05; Fig. 2A and Supplementary Table 1). The lateral occipital cortex showed decreased CT in patients compared to controls in both the left hemisphere and right hemisphere. The left caudal anterior cingulate, right isthmus cingulate, and right pericalcarine also showed decreased CT in patients. Cortical regions with increased CT in patients included the pars opercularis from both the left hemisphere and right hemisphere, the right rostral middle frontal cortex, right temporal pole, and right superior temporal cortex. In general, we observed more decreased CT (atrophy) in caudal regions of the cortex compared to rostral regions that showed increased CT (hypertrophy).

**Figure 2.**
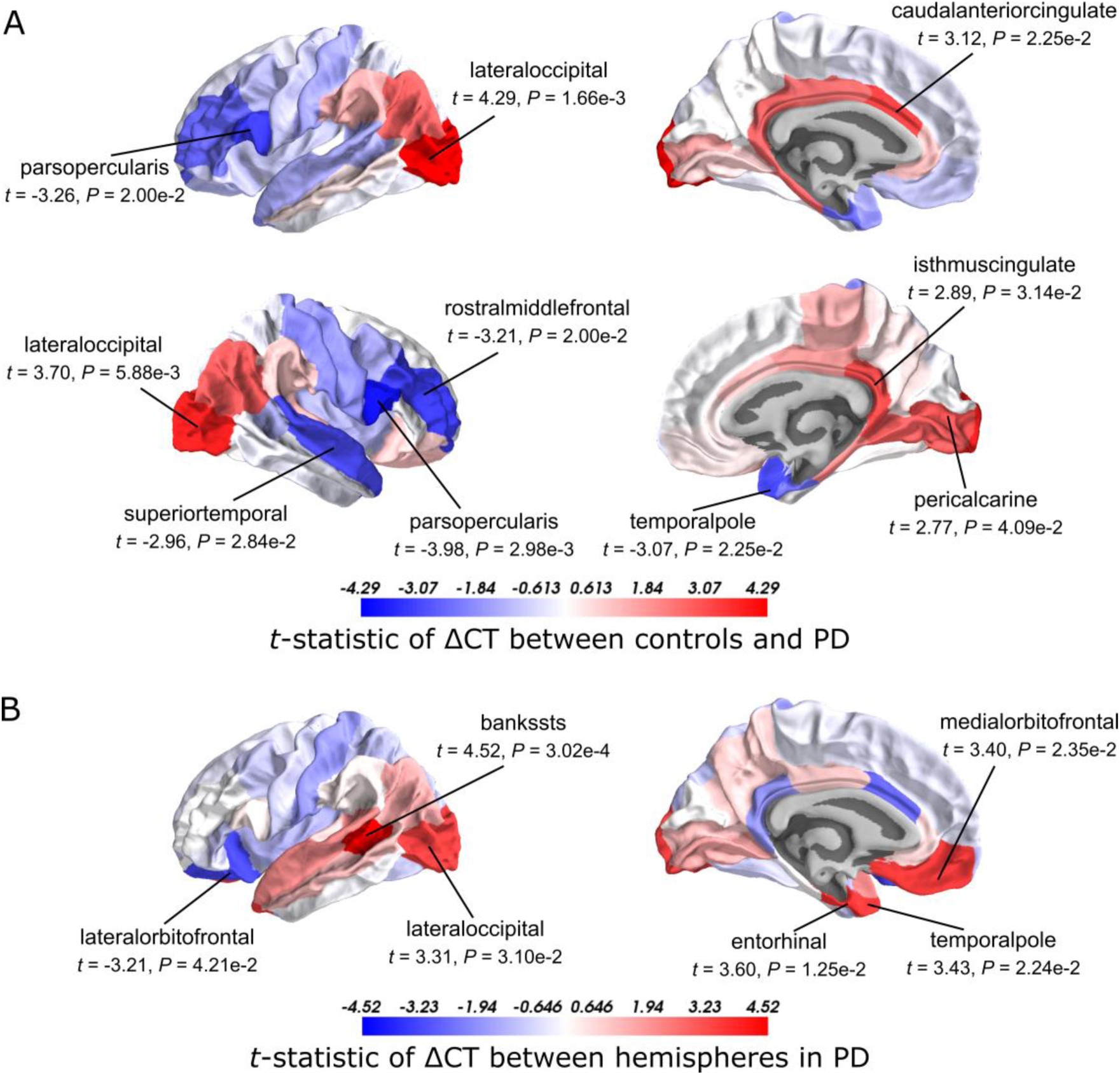
*t*-Statistics of cortical thickness changes (ΔCT) across cortical regions. (A) CT was assessed between Parkinson’s disease patients and healthy controls. Higher *t*-statistics (red) indicate a larger CT in controls compared to the CT in patients and thus corresponds to cortical atrophy. (B) CT in the left hemisphere compared to the right hemisphere in Parkinson’s disease patients. Higher *t*-statistics (red) indicate a larger CT in the right hemisphere compared to the left hemisphere and thus corresponds to cortical atrophy in the left hemisphere. *P*-values are BH-corrected and significant regions (*P* < 0.05) are labeled.

### CT changes between hemispheres in Parkinson’s disease

Clinical symptoms appear asymmetrical at disease onset with the left hemisphere being more susceptible to degeneration than the right (Claassen *et al*., 2016). To assess whether this asymmetry is reflected also in the observed atrophy patterns, we compared the CT between the left and right hemisphere for each of the 34 cortical regions in Parkinson’s disease patients. We found six cortical regions that showed significant hemispheric differences (BH-corrected *P* < 0.05; Fig. 2B and Supplementary Table 2). For five out of six significant regions, CT was indeed smaller in the left hemisphere compared to the right: banks of superior temporal sulcus, entorhinal cortex, temporal pole, medial orbitofrontal cortex, and lateral occipital cortex. For the lateral orbitofrontal cortex, the CT was larger in the left hemisphere compared to the right.

### Gene expression patterns predictive of CT changes in Parkinson’s disease patients

To identify the molecular mechanisms underlying CT changes in Parkinson’s disease, we integrated the imaging features with brain-wide gene expression profiles from the AHBA (Fig. 1C). Using PLS *model-1* (see Methods), the expression of all 20,017 genes in 34 brain regions from the left hemisphere was used as predictor variables and we used the *t*-statistics of ΔCT between Parkinson’s disease patients and controls in the 34 regions (Supplementary Table 1) as a single response variable. The number of AHBA samples varied between 0 and 92 for each one of the six brain donors and 34 cortical regions (Supplementary Table 3).

The PLS components that explain maximum covariance between the input space and the response variable are derived from successively deflated predictor and response matrices. Hence, the first component of the predictor matrix, *component-1*, has maximum covariance with the first component of the response matrix, and the second component of the predictor matrix, *component-2*, has maximum covariance with the second component of the response matrix, etc. Since PLS *model-1* has a single response variable, *component-1* of the response matrix is equal to a scaled version of the single response variable. As such, we only examined PLS *component-1* of the predictor matrix (additional checking with leave-one-out cross-validation showed that the optimal number of components is indeed one, Supplementary Fig. 2).

The scores of PLS *component-1* of the predictor variables (genes) showed a caudal-to-rostral expression pattern (Fig. 3A) that was correlated with CT changes in Parkinson’s disease brains (Fig. 3B), i.e. gene expression of PLS *component-1* was high in caudal regions associated with atrophy and low in regions associated with hypertrophy. The Pearson correlation between the PLS *component-1* scores of the predictor variables (gene expression) and the response variable (*t*-statistics of ΔCT) was 0.58, and explained 20.5% of the variance in gene expression and 34.2% of the variance in CT changes. Cortical atrophy was highest in the lateral occipital cortex and related to high PLS *component-1* scores. The pericalcarine showed the highest PLS *component-1* score. These results showed that the expression profiles of a weighted combination of genes can be predictive of CT changes in Parkinson’s disease.

**Figure 3.**
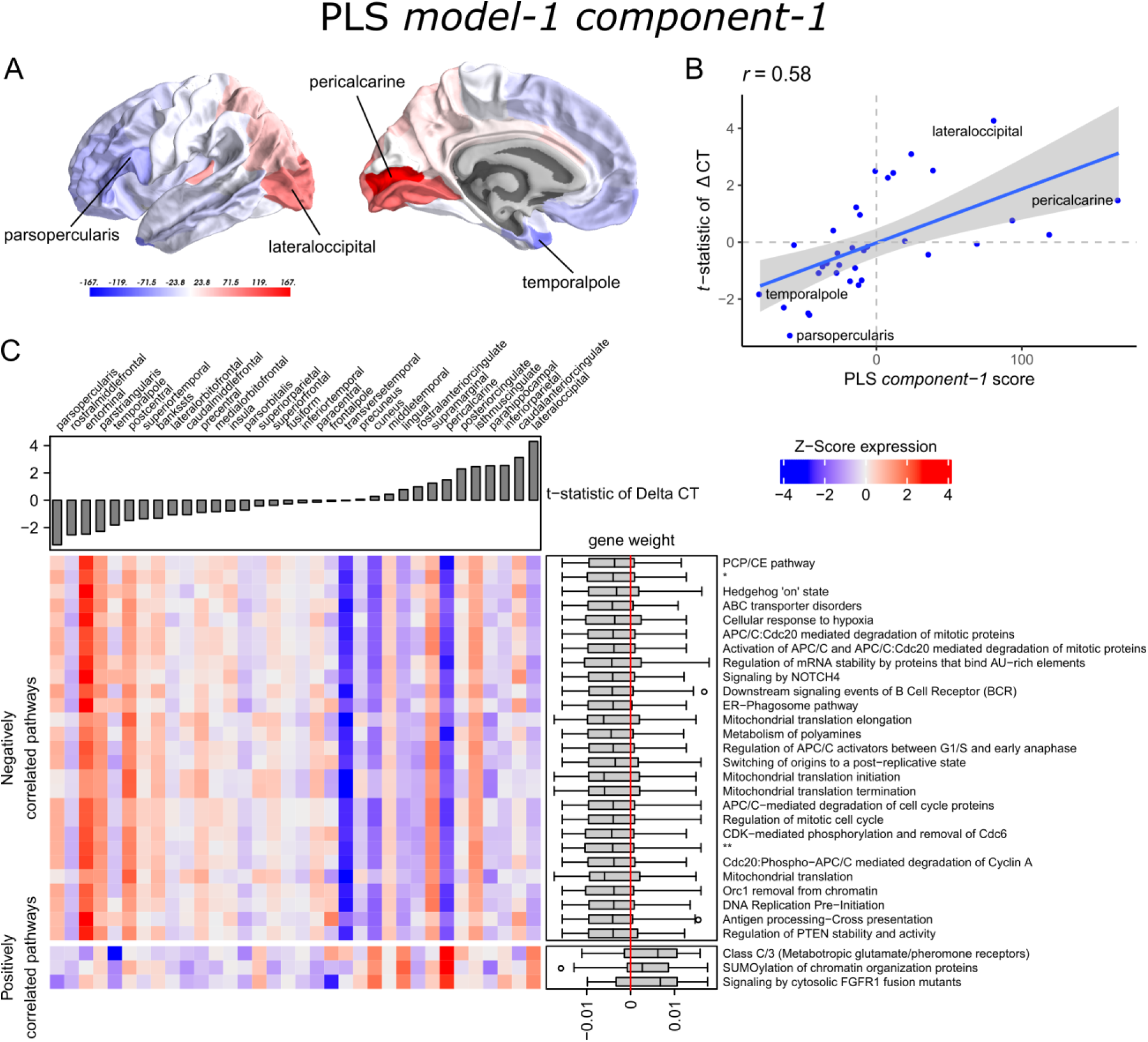
Transcriptional signatures to predict *t*-statistics of ΔCT between Parkinson’s disease patients and controls in PLS *model-1*. (A) PLS *component-1* scores of predictor variables (gene expression) visualized in cortical regions (lateral and medial view of the left hemisphere). (B) Regression fit of the latent predictor variable, PLS *component-1* scores, with the single response variable, CT changes in Parkinson’s disease measured as the *t*-statistics of ΔCT between Parkinson’s disease patients (149) and controls (369) across the 34 cortical regions. (C) Mean expression of genes in the top 30 significant pathways (rows) across cortical regions (columns). A complete heatmap with all significant pathways is given in Supplementary Fig. 4. The correlation between transcriptomic signatures and CT changes in Parkinson’s disease across cortical regions is predicted by the gene weights for PLS *component-1* shown in boxplots for each pathway where the median weight is either negative or positive. Negatively correlated pathways show high gene expression in regions with low *t*-statistics of ΔCT and gene expression decreases in regions with higher *t*-statistics of ΔCT. In our analysis, negative *t*-statistics correspond to increased CT (cortical hypertrophy) and positive *t*-statistics of ΔCT correspond to decreased CT (cortical atrophy). Positively correlated pathways show low expression in regions with low *t*-statistics of ΔCT and expression increases in regions with higher *t*-statistics of ΔCT. * = APC:Cdc20 mediated degradation of cell cycle proteins prior to satisfation of the cell cycle checkpoint. ** = APC/C:Cdh1 mediated degradation of Cdc20 and other APC/C:Cdh1 targeted proteins in late mitosis/early G1.

### Functionality of genes predictive of CT changes

A PLS component of the predictor variables is a linear combination of weighted gene expression. We used the gene weights of PLS *component-*1 to perform GSEA analysis and revealed significant enrichment of 90 pathways, which were among others involved in DNA damage checkpoints, stabilization of p53, regulation of apoptosis, mitochondrial translation, and SUMOylation of chromatin organization proteins (Supplementary Table 4). High overlap of genes between the enriched pathways suggested that these functional processes are highly related to each other (Supplementary Fig. 3).

Significant pathways are either positively or negatively correlated with CT changes based on the median weight of genes within pathways. Out of the 90 pathways that were significantly enriched, three pathways were positively correlated with the *t*-statistic of ΔCT. These included SUMOylation of chromatin organization proteins, signaling by cytosolic FGFR1 fusion mutants, and class C/3 (Metabotropic glutamate/pheromone receptors). Higher mean expression of genes within these three pathways is related to cortical atrophy (higher *t*-statistics of ΔCT); as apparent in the lateral occipital cortex (Fig. 3C and Supplementary Fig. 4). The positive correlation also indicates that a lower expression of these pathways is related to cortical hypertrophy (lower *t*-statistics of ΔCT). We found 87 negatively correlated pathways (median gene weight < 0). These pathways seem to play a role in the mitochondrial regulation of mitosis as we found pathways for mitochondrial translation, the regulation of mitotic cell cycle, p53-(in)dependent DNA damage checkpoints, and the degradation of mitotic proteins, such as cyclins A, and D. In general, the mean expression of genes in the negatively correlated pathways was high in cortical regions that showed hypertrophy, such as the pars opercularis or the entorhinal cortex.

### Relationships between clinical scores and cortical thickness

Next, we set to understand the relationship between CT in 34 cortical regions and clinical scores of Parkinson’s disease patients. Linear regression was used to predict clinical scores from CT across patients and obtain regression coefficients, *β*_1_, for each cortical region and clinical domain (Equation 1). We assessed the *t*-statistics of the regression coefficients instead of the coefficients *β*_1_ themselves (H_0_: *β*_1_ = 0) (Fig. 4). Negative *t*-statistics showed that most combinations of cortical regions and clinical features are negatively correlated. For all clinical features, higher scores also indicate more severe symptoms, except for MMSE scores where lower scores indicate more severe symptoms, and thus showed positive relationships with CT. In most regions, age at onset (AGEONSET) also showed positive relationships with CT, indicating that age at onset has an effect on the loss of CT. While these general interpretations apply to most cortical regions, some regions showed different relationships with CT. For example, CT in the rostral anterior cingulate is negatively related to age at onset, and positively related to MDS-UPDRS 4 scores.

**Figure 4.**
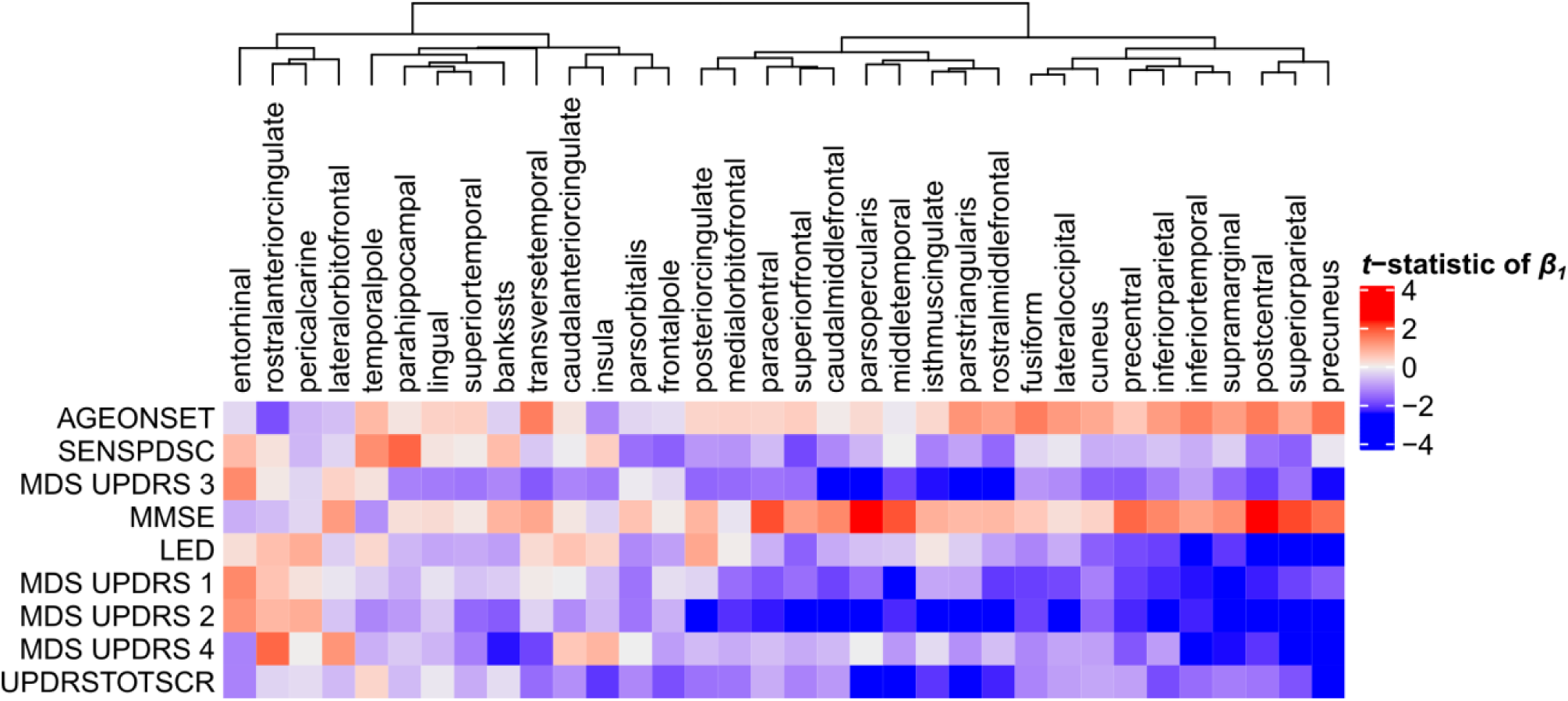
Relationship between clinical scores and CT across Parkinson’s disease patients. Linear regression was used to predict clinical scores from CT across at most 123 Parkinson’s disease patients (Supplementary Fig. 1). Separate models were used for each clinical feature (row) and cortical region (column) to obtain regression coefficients, see Equation 1. The heatmap shows the two-sided *t*-statistics of the regression coefficient when tested for H_0_: ***β***_**1**_ = 0. Regions (columns) are clustered based on complete linkage of the Euclidean distance of the *t*-statistics of ***β***_**1**_.

### Genes predictive of relationships between clinical scores and cortical thickness

With PLS *model-2*, we examined gene expression patterns that are predictive of the relationship between CT and clinical scores measured as *t*-statistics of the correlation coefficients *β*_**1**_ in Equation 1 (Fig. 1C). We selected the first two PLS components for further analysis, which explained 36% of the variance of the predictor variables and 37% of the variance of the response variables (Supplementary Fig. 5). PLS *component-1* scores of the predictor variables showed a ventral-to-dorsal gene expression pattern (Fig. 5A) that is correlated with the PLS *component-1* scores of the response variables (Pearson *r* = 0.76, Fig. 5B). The dorsal regions include the postcentral gyrus which is part of the primary somatosensory cortex. PLS *component-2* scores of the predictor variables showed a caudal-to-rostral gene expression pattern (Pearson *r* = 0.56, Fig. 6A) that is correlated with the PLS *component-2* scores of the response variables (Fig. 6B). Moreover, we assessed PLS *component-3* (Pearson *r* = 0.76 between the predictors and response variables), which additionally explained 9% variance of the predictor variables and 11% variance of the response variables. However, further analysis revealed there were no enriched pathways for *component-3* limiting the functional interpretation of this component.

**Figure 5.**
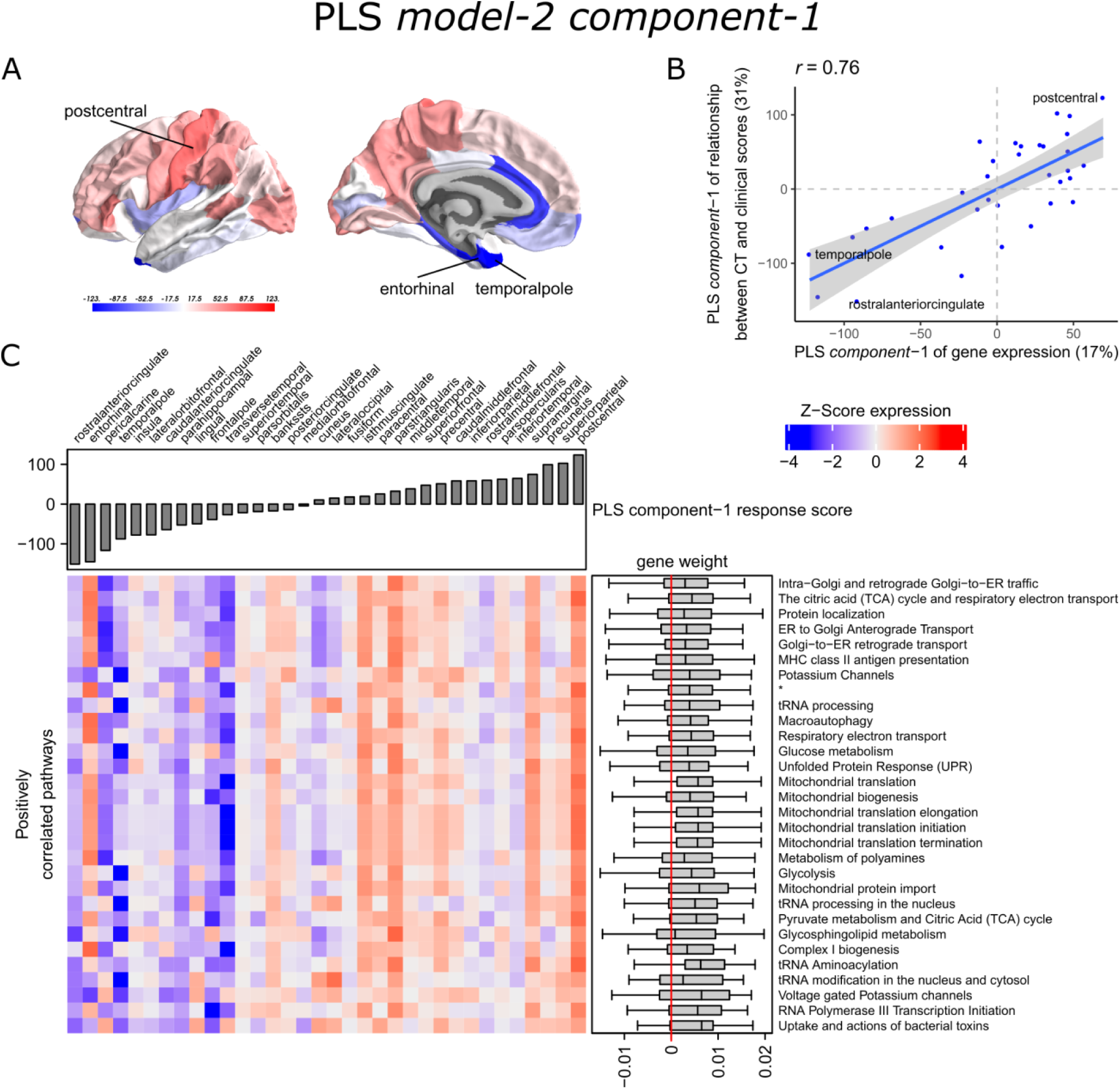
Transcriptional signatures of PLS *component-1* in PLS *model-*2 predictive of the relationship between cortical thickness (CT) and clinical scores. (A) PLS scores for PLS *component-1* of the predictor variables (gene expression) and (B) its correlation with PLS *component-1* of the response variables (*t*-statistic of ***β***_**1**_ in Equation 1). Axes show the percentage of explained variance for each component; *r* indicates the Pearson correlation. (C) Mean expression across cortical regions (columns) of genes in the top 30 significant pathways (rows). A complete heatmap with all significant pathways is given in Supplementary Fig. 8. * = Respiratory electron transport, ATP synthesis by chemiosmotic coupling, and heat production by uncoupling proteins.

**Figure 6.**
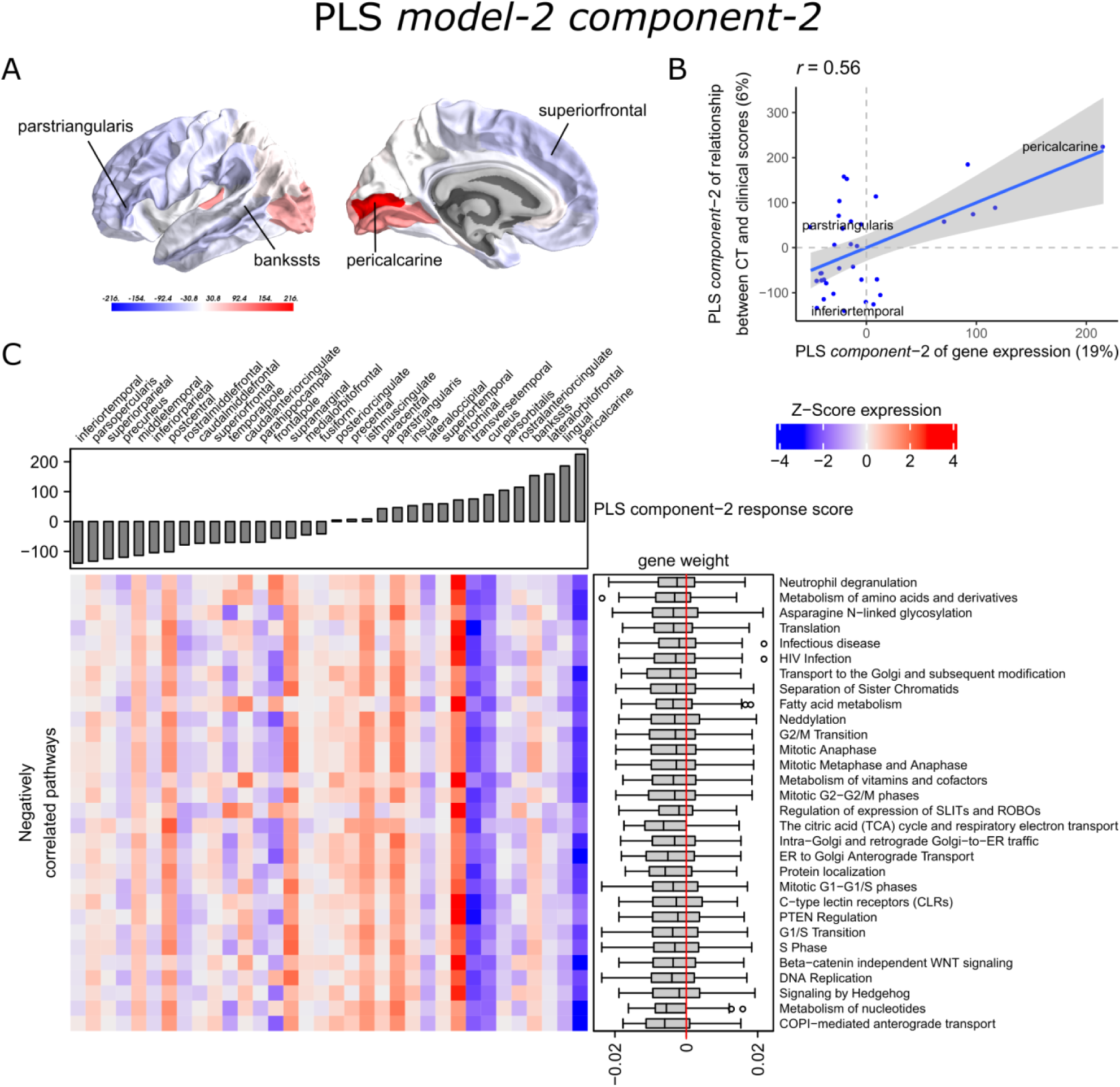
Transcriptional signatures of PLS *component-2* in PLS *model-2* predictive of the relationship between cortical thickness (CT) and clinical scores. (A) PLS scores for PLS *component-2* of the predictor variables (gene expression) and (B) its correlation with PLS *component-1* of the response variables (*t*-statistic of ***β***_**1**_ in Equation 1). Axes show the percentage of explained variance for each component; *r* indicates the Pearson correlation. (C) Mean expression across cortical regions (columns) of genes in the top 30 significant pathways (rows). A complete heatmap with all significant pathways is given in Supplementary Fig. 9.

PLS *component-1* and *component-2* of the predictor variables showed 144 and 230 significantly enriched pathways, respectively, with 54 overlapping pathways between the two components (Supplementary Table 5 and 6). Both components showed a cluster of related pathways involved in anterograde and retrograde transport between Golgi and endoplasmic reticulum (ER), and asparagine N-linked glycosylation (Supplementary Fig. 6 and 7). Other pathways that overlapped between the two components included macroautophagy, mitochondrial translation, mitochondrial biogenesis, mitochondrial protein import, DNA damage/telomere stress induced senescence, oxidative stress induced senescence, and protein localization.

Furthermore, PLS *component-1* showed enrichment of pathways involved in tRNA and rRNA processing in the nucleus and mitochondrion, voltage-gated potassium channels, uptake and actions of bacterial toxins, and interleukin signaling. PLS *component-2* showed strong enrichment of neutrophil degranulation, DNA replication, p53-(in)dependent DNA damage response, and chaperonin-mediated protein folding, and tubulin folding. Notably, the gene expression pattern of PLS *component-2* was also associated with several sumoylation pathways and pathways involved in mitotic cell cycles and the degradation of mitotic proteins (Supplementary Table 6).

The enriched pathways for PLS *component-1 and component-2* either showed negative or positive median gene weights that inform about the sign of the correlation between genes within a pathway and the PLS component score of the response variables (Fig. 5C, Fig. 6C, Supplementary Fig. 8 and 9). For example, the expression of genes within pathways relating to mitochondrial processes increases for higher PLS *component-1* scores of the response variables. We further assessed PLS *component-1 and component-2* scores of the predictor variables and their correlation with each individual response variable, which are the clinical features and their relationship with CT in Parkinson’s disease patients (Fig. 7). The rostral-to-dorsal expression pattern of PLS *component-1* is highly predictive of the relationship between CT and MMSE score in patients (Pearson’s *r* = 0.71). Thus, pathways associated with PLS *component-1* may play an important role in cognitive circuits, which seems to be apparent based on their expression in the postcentral gyrus, but also the entorhinal cortex. PLS *component-2* scores showed low correlations with the clinical features and their relation with CT across cortical regions, and suggests weak associations between the expression patterns of PLS *component-2* and the response variables.

**Figure 7.**
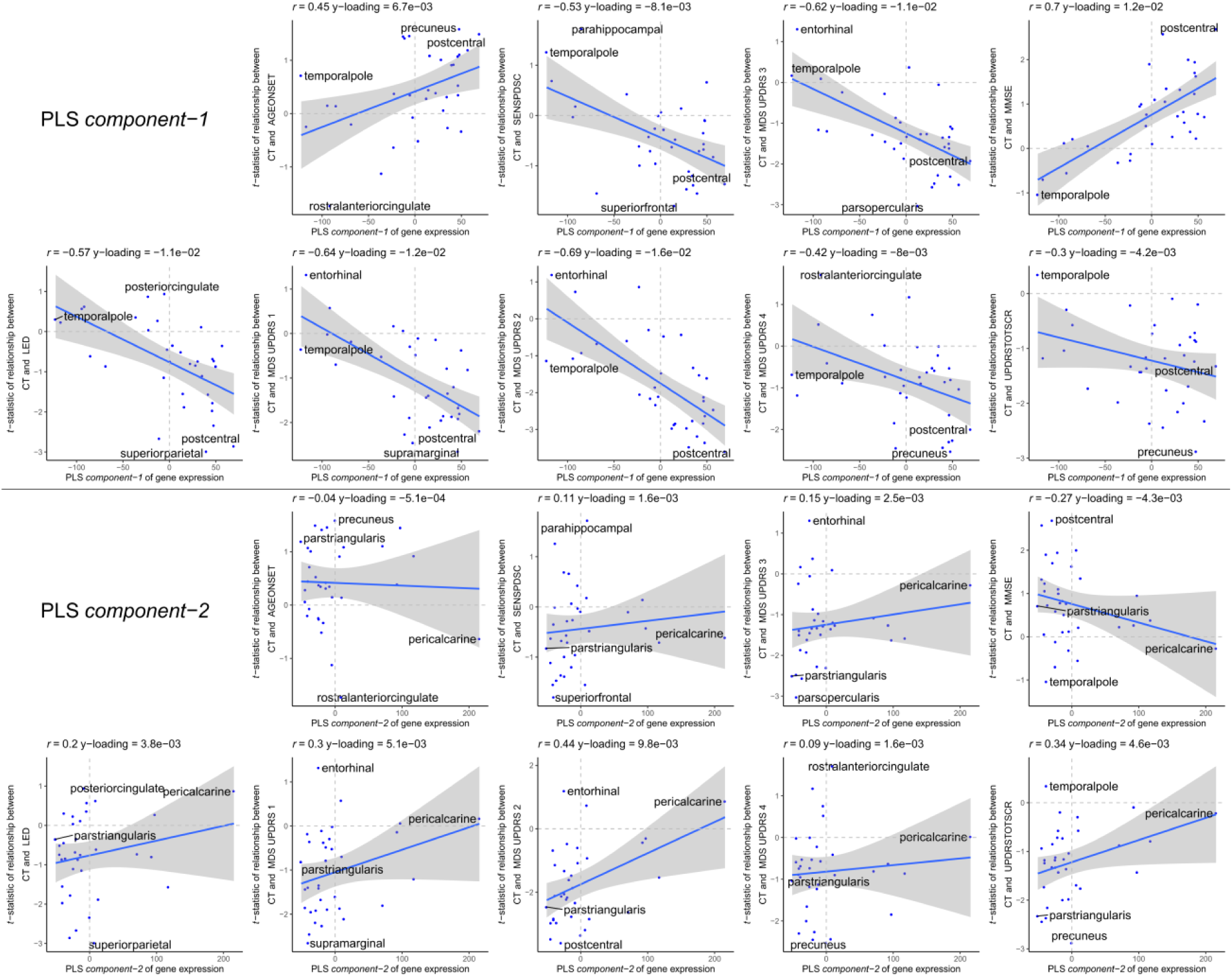
Correlations between PLS *model-2 component-1 and component-2* scores of the predictor variables and individual response variables. Each plot shows the correlation between the predictor variables of gene expression (x-axis) and the response variables which are the relationships between CT and scores of a clinical feature across cortical regions (y-axis). On top of each plot, the Pearson correlation and the Y-loadings (Q in Eq. 4 and 8) are shown; both values tell something about the sign (-/+) and magnitude (high/low) of the correlation. Each point or sample is one of the 34 cortical regions. Regions are labeled for those with minimum or maximum value along one of the axes.

## Discussion

We found a caudal-to-rostral gene expression pattern that was correlated with CT changes in Parkinson’s disease (PLS *model-1*); cortical atrophy was found in caudal regions while rostral regions showed cortical hypertrophy. This transcriptional signature was highly enriched for genes in biological pathways associated with mitochondrial translation and mitotic cell cycle regulation. We also found a ventral-to-dorsal and caudal-to-rostral gene expression pattern that was correlated with the relationship between CT and clinical domains of Parkinson’s disease (PLS *model-2*). Both transcriptional signatures were associated with similar pathways, including macroautophagy and Golgi-ER trafficking, and may be involved in the effect of CT on clinical scores, namely MMSE scores for cognitive assessment.

The CT analyses between disease conditions and hemispheres in patients revealed cortical regions that are susceptible to atrophy. Cortical atrophy in Parkinson’s disease commonly occurs asymmetrical, with a preference for the left hemisphere, particularly in the early disease stages (Brück *et al*., 2004; Mak *et al*., 2014; Pereira *et al*., 2014; Claassen *et al*., 2016). Here, we showed that five out of six regions with significant CT changes between hemispheres, indeed revealed more atrophy in the left hemisphere. Two cortical regions that showed significant changes between patients and controls, also showed changes between the left and right hemisphere. Our findings are in line with those of a previous study showing that cortical atrophy in Parkinson’s disease most prominently affects the lateral occipital cortex, particularly in the left hemisphere (Freeze *et al*., 2018). The temporal pole showed hypertrophy in patients compared to controls, which was only significant in the right hemisphere. However, our analysis between hemispheres of Parkinson’s disease brains suggests that the left temporal pole is more susceptible to CT loss than the right hemisphere. The remaining regions that were susceptible to CT changes showed atrophy in either the left or right hemisphere; however differences between hemispheres in patients could not be confirmed. All 10 regions that were different between patients and controls, except the pericalcarine, were earlier identified as part of two structural covariance networks that were related to gray matter atrophy in the same Parkinson’s disease dataset as in this study (de Schipper *et al*., 2017). Overall, we observed atrophy in caudal regions, which earlier has been associated with late stage Parkinson’s disease (Claassen *et al*., 2016).

With our findings of the PLS models we interpret gene expression patterns of the healthy brain in relation to imaging features observed in Parkinson’s disease. The six adult donors of the AHBA had no known neuropsychiatric or neuropathological history (Hawrylycz *et al*., 2012), however it is unknown whether these individuals could have developed neurodegenerative diseases later in life. The observed spatial gene expression patterns reflect the physiological conditions in the adult healthy brain and are informative of important molecular mechanisms that are vulnerable in Parkinson’s disease. The biological pathways found for PLS *model-1* were closely related as they shared many similar genes. These interrelated pathways suggest a strong functional relationship between molecular processes involving mitotic cell cycle, mitochondrial translation, transport between ER and Golgi, DNA damage checkpoints, and sumoylation. We found that differential regulation of these molecular processes across the brain was associated with CT changes observed in Parkinson’s disease. Similar pathways were found in PLS *model-2* with multiple response variables corresponding to the relationships between CT and nine clinical domain scores in Parkinson’s disease.

There is evidence that impaired cell cycle control plays a role in the pathogenesis of neurodegenerative diseases. In healthy conditions, differentiated neuronal cells become quiescent cells that cannot re-enter the cell cycle, however in neurodegenerative diseases they are reactivated which is associated with increased cell death (Bonda *et al*., 2018). Cell cycle checkpoints are controlled by cyclins that guide the cell from one phase to the next phase and its expression can induce cell cycle re-initiation (Walton *et al*., 2019). Here, we found that regional expression of pathways associated with the degradation of cell cycle proteins in healthy conditions were negatively correlated with CT changes in Parkinson’s disease, i.e. higher expression was associated with cortical hypertrophy in rostral regions such as the pars opercularis and temporal pole. Reversely, we observed low expression of protein degradation pathways in caudal regions that were associated with atrophy, and therefore suggests that regions with low expression are more vulnerable to improper degradation of cell cycle proteins leading to cell cycle initiation. This indicates that regions with low expression of such essential pathways are predisposed to neurodegeneration.

We found that the expression of several pathways associated with DNA replication and p53-(in)dependent DNA damage responses and checkpoints were correlated with CT changes. DNA replication during the S-phase may control the survival of post-mitotic cells by DNA repair mechanisms or apoptosis followed by DNA damage, which seems to be the case in neurodegenerative diseases (Tokarz *et al*., 2016). Furthermore, DNA damage response signaling can be modulated by tumor suppressor p53 and may also contribute to apoptosis in aging and age-related neurodegenerative disorders (Mohammadzadeh *et al*., 2019). These pathways showed similar expression patterns as those associated with the mitotic cell cycle, and therefore a lower expression of these DNA damage response pathways in caudal regions is related to cortical atrophy in Parkinson’s disease.

Similar caudal-to-rostral expression patterns were found for pathways associated with mitochondrial translation. Increased risk for Parkinson’s disease has been associated with mutations in *SNCA, PARK2* (parkin), *PINK1, DJ-1*, and *LRRK2* which have been linked to mitochondrial function and oxidative stress (Yan *et al*., 2012). *PINK1* and parkin mediates clearance of damaged mitochondria by mitophagy and may therefore influence mitotic cell cycle progression (Sarraf *et al*., 2019). *PINK1* also regulates both retrograde and anterograde axonal transport of mitochondria via axonal microtubules (Liu *et al*., 2012) The interaction between *PINK1* and parkin is likely involved in mitochondrial quality control mechanisms, where anterograde transport of damaged mitochondria is reduced and retrograde transport is enhanced for elimination by mitophagy in the neuronal cell body (Lionaki *et al*., 2015).

A cluster of pathways involved in ER-Golgi traffic were found enriched for PLS *model-2 component-1 and component-2*, and involved both ER-to-Golgi anterograde and Golgi-to-ER retrograde transport. *Component-1* showed a ventral-to dorsal gene expression pattern that was associated with higher correlations between CT and clinical scores, namely, the mental state of Parkinson’s disease patients and the performance of motor functions. The pathways involved in ER-Golgi traffic were notably high expressed in the postcentral gyrus which contains the somatosensory cortex that is known for its role in processing sensory information and the regulation of emotion (Kropf *et al*., 2019). Our results suggest that genes in ER-Golgi traffic pathways are important for cognitive functions controlled by the postcentral gyrus. Genes involved in ER-Golgi vesicle trafficking have the ability to modify α-synuclein toxicity in yeast (Cooper *et al*., 2006). Moreover, fragmentation to the Golgi apparatus has been associated with the accumulation of aberrant proteins in neurodegenerative diseases, including α-synuclein (Fan *et al*., 2008). A study in yeast models has showed that α-synuclein expression modulates ER stress signaling response and inhibits viral infections and viral replication (Beatman *et al*., 2016). We found several pathways associated to HIV and influenza infections that were correlated to the relationship between CT and clinical scores. Another pathway that shared overlapping genes with those involved in ER-Golgi traffic was asparagine N-linked glycosylation, which is a biochemical linkage important for the structure and function of proteins. The *N*-glycosylated proteins are synthesized essentially in the ER and Golgi through sequential reactions and aberrant glycolysation of proteins may lead to inflammation and mitochondrial dysfunction in Parkinson’s disease and consequently to a cellular overload of dysfunctional proteins (Videira and Castro-Caldas, 2018).

We found that the expression of genes involved in sumoylation of chromatin organization proteins was correlated with CT changes, i.e. higher expression within caudal brain regions, such as the pericalcarine and the lateral occipital cortex, was associated with greater atrophy in Parkinson’s disease. Therefore, higher activity of sumoylation events may play a role in the regional vulnerability to neurodegeneration observed in Parkinson’s disease. On the other hand, lower expression of these pathways, such as in the pars opercularis, was associated with hypertrophy in rostral regions, suggesting that lower expression of sumoylation pathways has a protective effect. Additionally, the higher expression of sumoylation pathways was associated with higher correlations between CT and clinical scores as projected by PLS *component-2* in *model-2*. Sumoylation involves small ubiquitin-like modifier (SUMO) proteins that increase in response to cellular stress, such as DNA damage and oxidative stress, and can promote α-synuclein aggregation and Lewy body formation (Bologna and Ferrari, 2013; Eckermann, 2013; Rott *et al*., 2017). Several proteins associated with inherited forms of Parkinson’s disease are targets modified by SUMO regulating mitochondrial processes, these include α-synuclein, DJ-1, and parkin (Guerra de Souza *et al*., 2016). Sumoylation has been associated with several diseases, including cancers, cardiac diseases, and neurodegenerative diseases (Yang *et al*., 2017). In cancer, sumoylation mediates cell cycle progression and plays an essential role during mitosis (Eifler and Vertegaal, 2015). SUMO seems to promote cell death mediated by the p53 tumor suppressor protein, which may be responsible for the cell death of dopaminergic neurons in Parkinson’s disease (Eckermann, 2013). Our findings are in support of these hypotheses, and further suggest that sumoylation is important in specific cortical regions that are atrophic in Parkinson’s disease, such as the lateral occipital cortex.

Spatial gene expression data from Parkinson’s disease brains are limited in the number of brain donors and brain regions, which is mainly due to the limited availability of well-defined post-mortem Parkinson’s disease patients. Therefore, we used healthy gene expression from the AHBA to perform unbiased whole brain and whole transcriptome analysis. Gene expression for all the six healthy adult donors in AHBA was only available for the left hemisphere. Therefore, this study was restricted to the analysis of the left hemisphere when combining gene expression with MRI data. Furthermore, it is generally assumed that gene expression changes with age, however due to the limited number of brain donors in the AHBA, age-related differences in gene expression were not taken into account. In addition, MRI data from the patient and control groups were collected in different cohorts where different MRI scanners were used. However, both datasets were processed separately to obtain CT measurements per region. Thus, these morphological features could be directly compared between the two groups. Finally, to determine whether genes and pathways truly have predictive power of imaging features, both PLS models need to be validated with independent imaging cohorts of Parkinson’s disease.

We set out to find biological explanations for the selective regional vulnerability in Parkinson’s disease by correlating healthy gene expression in cortical regions with CT changes in Parkinson’s disease observed as atrophy and hypertrophy in neuroimaging data. We found genes that point towards pathways involved in cellular maintenance mechanisms that are well known in Parkinson’s disease and other neurodegenerative diseases, but were shown to be differently regulated across the brain. Sumoylation pathways showed opposite expression patterns across the brain compared to pathways associated with the regulation of mitotic cell cycle, p53-(in)dependent DNA damage response, mitochondrial translation, and ER-Golgi trafficking (Fig. 8). Nevertheless, all the enriched pathways were highly interconnected as shown by the number of shared genes and suggest a balanced interplay between sumoylation events and the other molecular mechanisms that seem to be important in controlling CT in different cortical regions. Moreover, we propose that dysfunctions of these pathways may impair motor and cognitive functions in Parkinson’s disease as a consequence of cortical atrophy.

**Figure 8.**
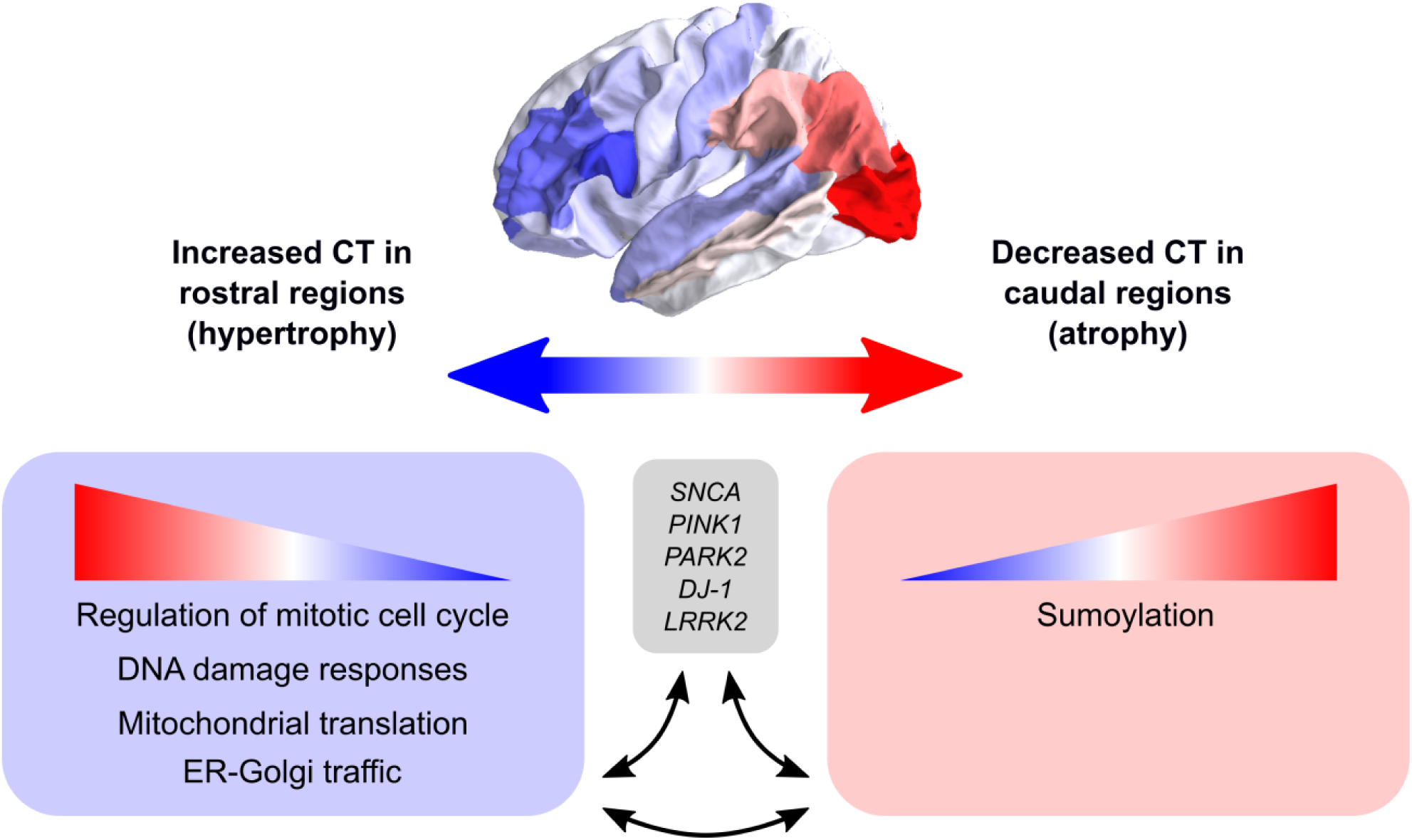
Schematic overview of the balance between biological pathways and their influence on CT across cortical brain regions. The big arrow indicates the caudal-to-rostral (red-to-blue) or rostral-to-caudal (blue-to-red) change in CT across cortical brain regions of Parkinson’s disease patients with red indicating decreased CT (atrophy) in caudal regions and blue indicating increased CT (hypertrophy) in rostral regions. Genes within pathways associated with sumoylation showed that the expression of these genes within the pathways increases from rostral to caudal regions. Other biological pathways that were correlated with CT changes in Parkinson’s disease included regulation of mitotic cell cycle, mitochondrial translation, DNA damage responses, and ER-Golgi traffic, and the involved genes showed decreasing expression patterns from rostral to caudal regions (or increasing from caudal to rostral regions). All enriched pathways shared many common genes and were generally associated with cellular maintenance mechanisms. Literature studies suggest that these biological pathways may be involved in the pathobiology of Parkinson’s disease through their interaction with genetic risk variants.

## Abbreviations

AHBA: Allen Human Brain Atlas;
CT: cortical thickness;
LED: levodopa equivalent dose;
MDS-UPDRS: Movement Disorder Society-sponsored revision of the unified Parkinson’s disease rating scale;
MMSE: mini-mental state examination;
PLS: partial least squares;
SENS-PD: severity of non-dopaminergic symptoms in Parkinson’s disease

## Acknowledgements

We would like to thank Laura J. de Schipper for discussions on atrophy patterns.

