## Supplemental Information for "Transcriptomic signatures associated with regional cortical thickness changes in Parkinson’s disease"

#### Supplementary Methods

##### Transcriptomic data

To filter and map probes to genes in the AHBA dataset, the data was concatenated across the six donors. We removed 10,521 probes with missing Entrez IDs, and 6,068 probes with low presence as they were expressed above the background in <1% of samples (PA-call containing presence/absence flag) (Hawrylycz *et al.*, 2015). The remaining 44,072 probes were mapped to 20,017 genes with unique Entrez IDs using the *collapseRows*-function in R-package WGCNA v1.64.1 (Langfelder and Horvath, 2008) as follows: i) if there is one probe, that one probe is chosen, ii) if there are two probes, the one with maximum variance across all samples is chosen (method="maxRowVariance"), iii) if there are more than two probes, the probe with the highest connectivity (summed adjacency) is chosen (connectivityBasedCollapsing=TRUE).

#### Partial least squares regression

As any linear regression method, PLS tries to find a relationship across $n$ samples:

$$Y = XB + B_0 \qquad (1)$$

, where data matrix $Y$ $(n \times p)$ with $p$ response variables is predicted by data matrix $X$ $(n \times m)$ with $m$ predictor variables, $B$ $(m \times p)$ is a matrix of regression coefficients, and $B_0$ $(n \times p)$ is a background term. Variables from both datasets $X$ and $Y$ are projected to latent variables such that they are maximally correlated:

$$X = TP^T + E \qquad (2)$$

$$Y = UQ^T + F \qquad (3)$$

, where $T$ And $U$ $(n \times c)$ are projections of $X$ and projections of $Y$, respectively, for $c$ components. $P$ $(m \times c)$ and $Q$ $(p \times c)$ are loading matrices, and $E$ and $F$ are the error terms. The scores for the latent variables $T$ and $U$ are optimized for maximal covariance between $X$ and $Y$:

$$U = TB + H \qquad (4)$$

, where $B$ $(m \times p)$ is a matrix of correlation coefficients, and $H$ $(n \times p)$ is the error term. Once the PLS model is calculated, the coefficients $B$ can be used to predict $Y$ directly from $X$ as in Equation 1, by using all $m$ variables in $X$ and all $p$ variables in $Y$ and their projections to $T$ and $U$, respectively. The estimation of each PLS component is determined iteratively, by successfully deflating $E$ and $F$, where projections of $X$ to the latent variables in $T$ are represented by $W$ $(m \times c)$. Since $W$ is calculated from deflated matrices, it cannot be used to represent weights that relate back to the original predictor matrix $X$. Instead, the weights can be represented by:

$$R = W(P^T W)^{-1} \qquad (5)$$

, where $R$ $(m \times c)$ relates back to the original matrix such that:

$$T = XR \qquad (6)$$

, and $R$ can be used to rank the predictor variables according to their contribution to $T$. Depending on the number of selected components, the regression coefficients $B$ are calculated by:

$$B = RQ^T \tag{7}$$

Based on the calculation of $B$ in Equation 7, we can also use $Q$ to assess the contribution of each response variable to the latent components in $U$.

#### Supplementary Figures

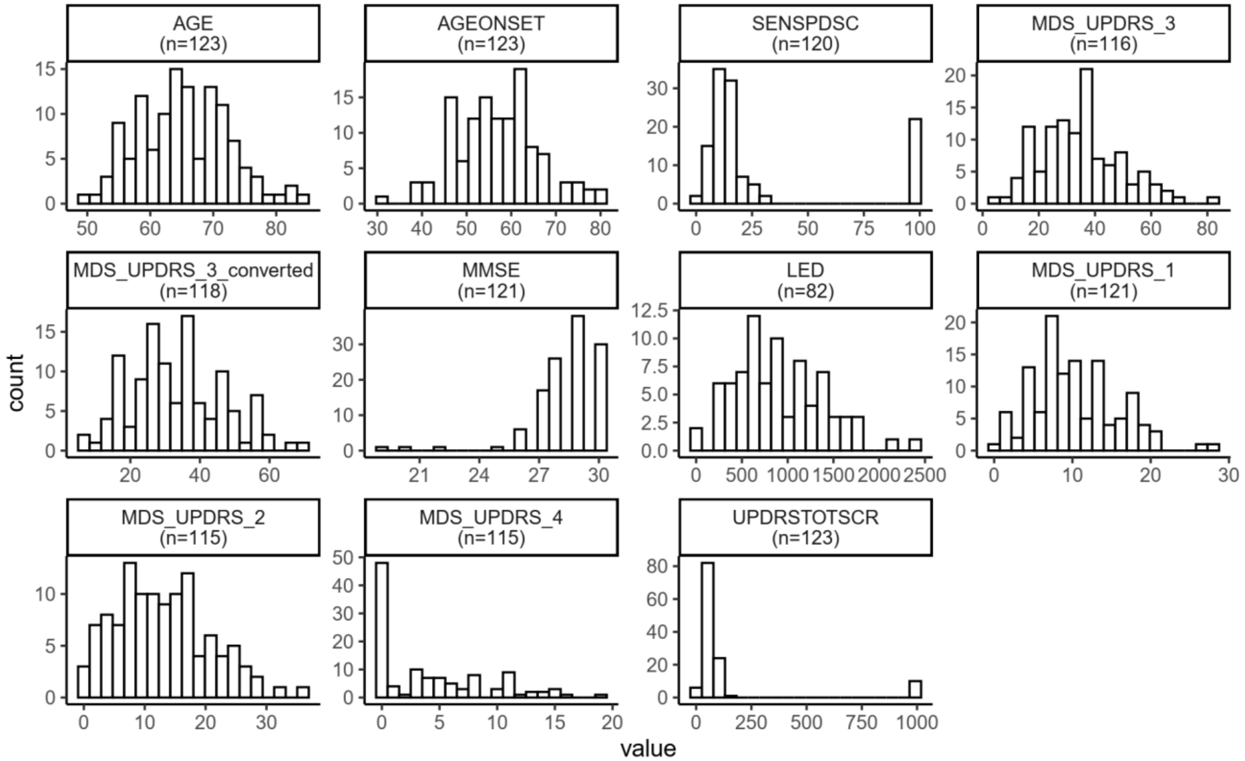


**Supplementary Figure 1 Score distribution for nine clinical domains of Parkinson's disease.** Scores are available for 82-123 patients.

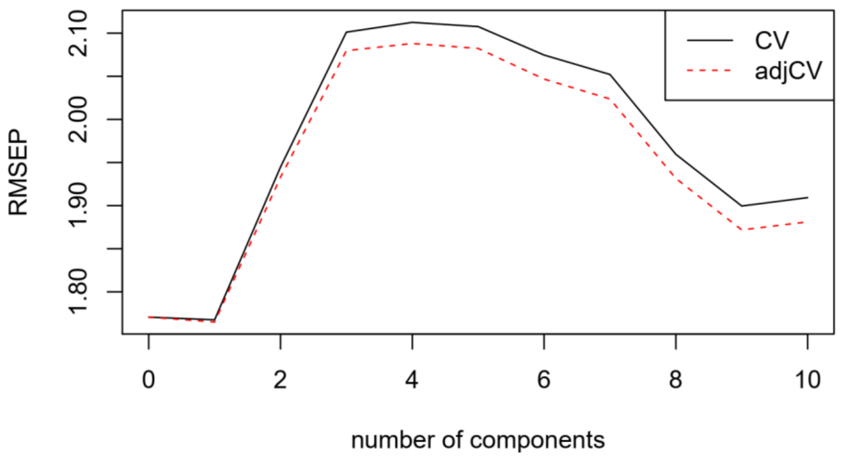


**Supplementary Figure 2 Root mean squared error of prediction (RMSEP) given the number of PLS components.**

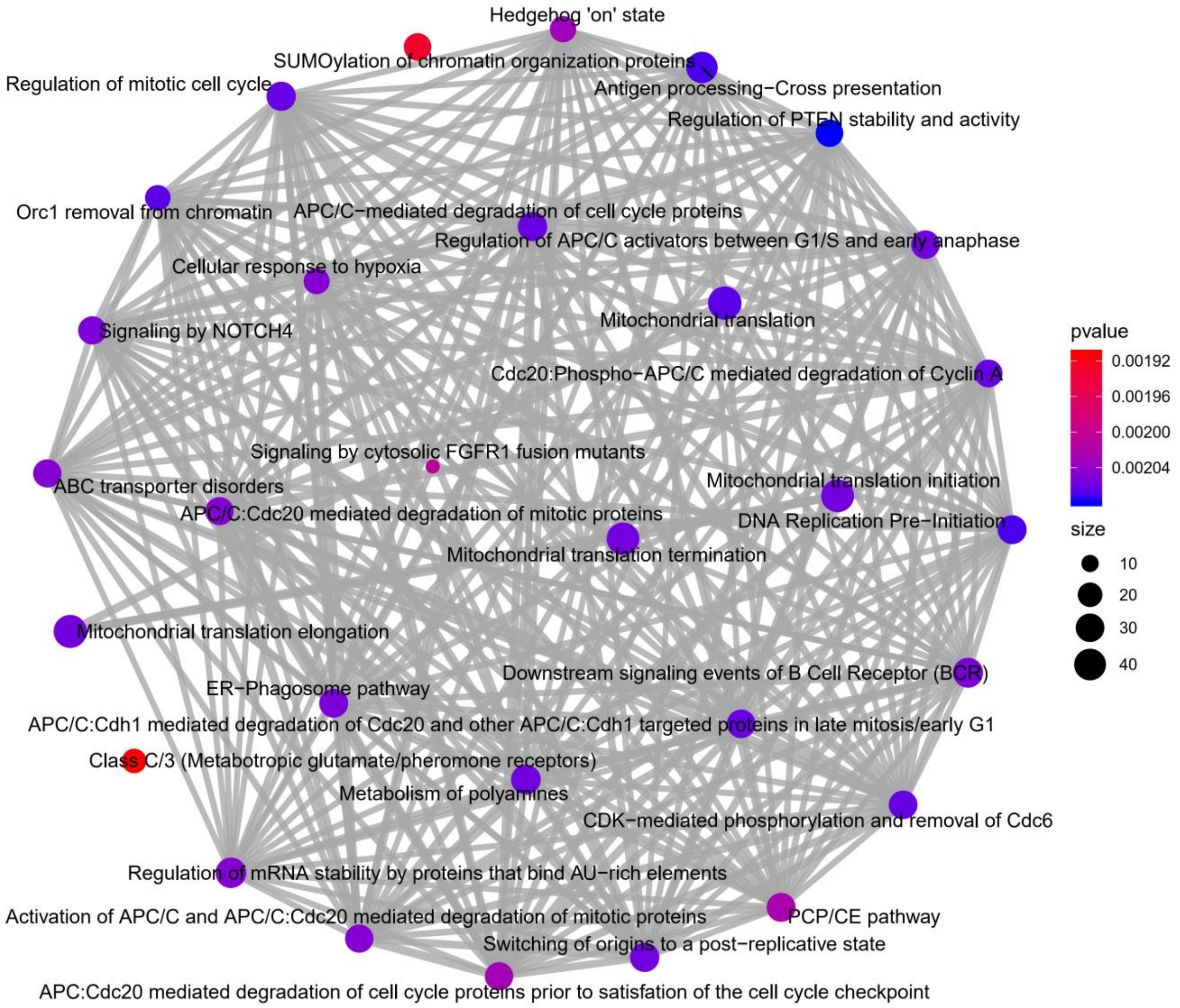

**Supplementary Figure 3 Pathway enrichment of genes ranked by their weights to PLS *component-1* of PLS *model-1*.** Enrichment was done using GSEA and Reactome pathways (BH-corrected $P < 0.05$). Node size indicates the number of genes in the pathway and node color shows the unadjusted *P*-values. Edge thickness indicates the number of overlapping genes between two pathways. This dense graph shows that the enriched pathways are highly related to each other. For visualization purpose, this graph only shows the top 30 (out of 90) enriched pathways. A list of all enriched pathways is given in Supplementary Table 4.

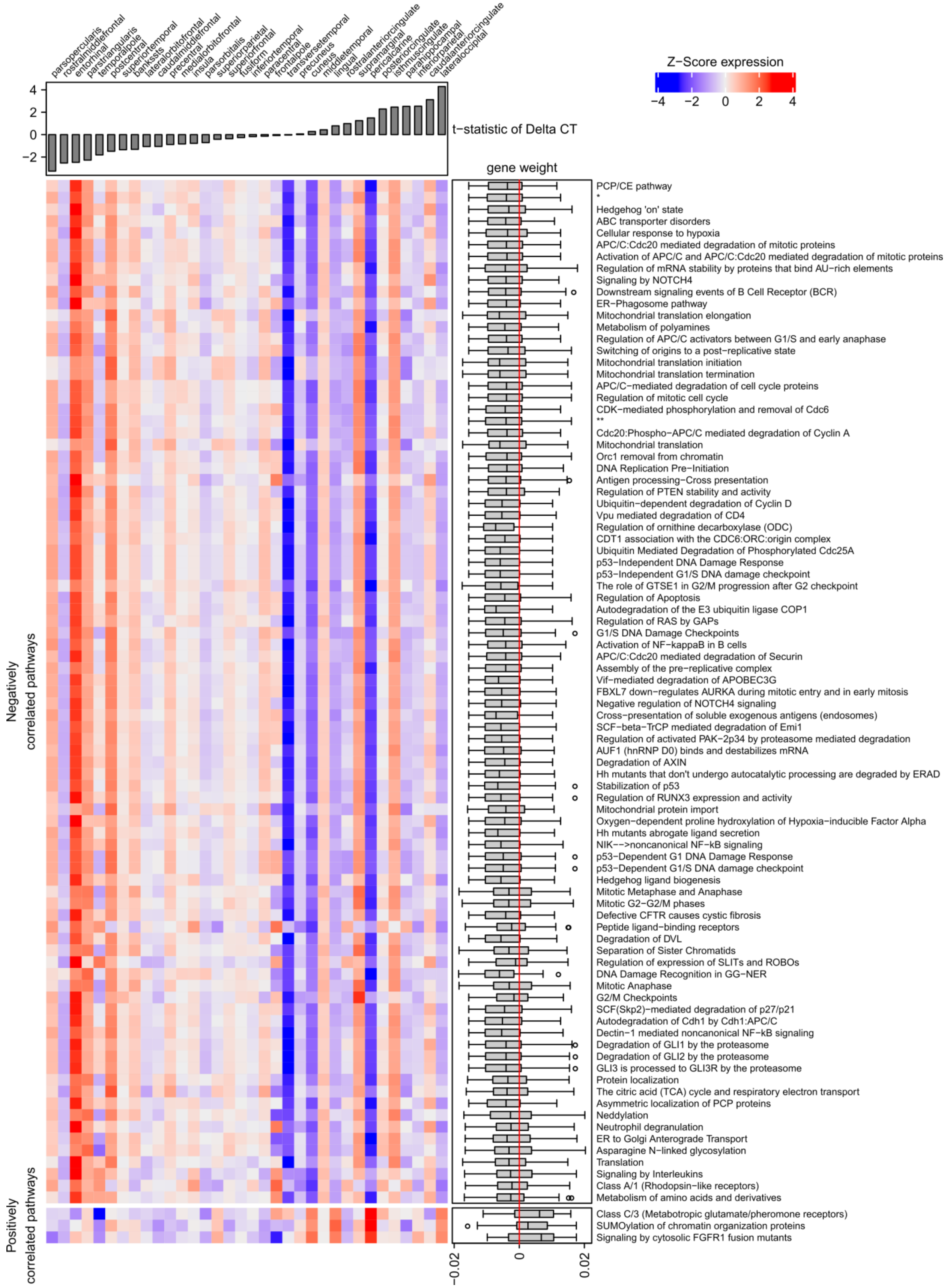
parsopercularis
rostralmiddlefrontal
entorhinal
parstriangularis
postcentral
temporalpole
superiortemporal
bankssts
lateralorbitofrontal
caudalmiddlefrontal
precentral
medialorbitofrontal
insula
parsorbitalis
superiorparietal
superiorfrontal
fusiform
inferiortemporal
paracentral
frontalpole
transversetemporal
precuneus
cuneus
middletemporal
lingual
rostralanteriorcingulate
supramarginal
pericalcarine
posteriorcingulate
isthmuscingulate
parahippocampal
inferiorparietal
caudalanteriorcingulate
lateraloccipital
Z−Score expression
−4
−2
0
2
4
4
2
0
−2
t−statistic of Delta CT
gene weight
Negatively correlated pathways
Positively correlated pathways
−0.02
0
0.02
PCP/CE pathway
*
Hedgehog 'on' state
ABC transporter disorders
Cellular response to hypoxia
APC/C:Cdc20 mediated degradation of mitotic proteins
Activation of APC/C and APC/C:Cdc20 mediated degradation of mitotic proteins
Regulation of mRNA stability by proteins that bind AU−rich elements
Signaling by NOTCH4
Downstream signaling events of B Cell Receptor (BCR)
ER−Phagosome pathway
Mitochondrial translation elongation
Metabolism of polyamines
Regulation of APC/C activators between G1/S and early anaphase
Switching of origins to a post−replicative state
Mitochondrial translation initiation
Mitochondrial translation termination
APC/C−mediated degradation of cell cycle proteins
Regulation of mitotic cell cycle
CDK−mediated phosphorylation and removal of Cdc6
**
Cdc20:Phospho−APC/C mediated degradation of Cyclin A
Mitochondrial translation
Orc1 removal from chromatin
DNA Replication Pre−Initiation
Antigen processing−Cross presentation
Regulation of PTEN stability and activity
Ubiquitin−dependent degradation of Cyclin D
Vpu mediated degradation of CD4
Regulation of ornithine decarboxylase (ODC)
CDT1 association with the CDC6:ORC:origin complex
Ubiquitin Mediated Degradation of Phosphorylated Cdc25A
p53−Independent DNA Damage Response
p53−Independent G1/S DNA damage checkpoint
The role of GTSE1 in G2/M progression after G2 checkpoint
Regulation of Apoptosis
Autodegradation of the E3 ubiquitin ligase COP1
Regulation of RAS by GAPs
G1/S DNA Damage Checkpoints
Activation of NF−kappaB in B cells
APC/C:Cdc20 mediated degradation of Securin
Assembly of the pre−replicative complex
Vif−mediated degradation of APOBEC3G
FBXL7 down−regulates AURKA during mitotic entry and in early mitosis
Negative regulation of NOTCH4 signaling
Cross−presentation of soluble exogenous antigens (endosomes)
SCF−beta−TrCP mediated degradation of Emi1
Regulation of activated PAK−2p34 by proteasome mediated degradation
AUF1 (hnRNP D0) binds and destabilizes mRNA
Degradation of AXIN
Hh mutants that don't undergo autocatalytic processing are degraded by ERAD
Stabilization of p53
Regulation of RUNX3 expression and activity
Mitochondrial protein import
Oxygen−dependent proline hydroxylation of Hypoxia−inducible Factor Alpha
Hh mutants abrogate ligand secretion
NIK−−>noncanonical NF−kB signaling
p53−Dependent G1 DNA Damage Response
p53−Dependent G1/S DNA damage checkpoint
Hedgehog ligand biogenesis
Mitotic Metaphase and Anaphase
Mitotic G2−G2/M phases
Defective CFTR causes cystic fibrosis
Peptide ligand−binding receptors
Degradation of DVL
Separation of Sister Chromatids
Regulation of expression of SLITs and ROBOs
DNA Damage Recognition in GG−NER
Mitotic Anaphase
G2/M Checkpoints
SCF(Skp2)−mediated degradation of p27/p21
Autodegradation of Cdh1 by Cdh1:APC/C
Dectin−1 mediated noncanonical NF−kB signaling
Degradation of GLI1 by the proteasome
Degradation of GLI2 by the proteasome
GLI3 is processed to GLI3R by the proteasome
Protein localization
The citric acid (TCA) cycle and respiratory electron transport
Asymmetric localization of PCP proteins
Neddylation
Neutrophil degranulation
ER to Golgi Anterograde Transport
Asparagine N−linked glycosylation
Translation
Signaling by Interleukins
Class A/1 (Rhodopsin−like receptors)
Metabolism of amino acids and derivatives
Class C/3 (Metabotropic glutamate/pheromone receptors)
SUMOylation of chromatin organization proteins
Signaling by cytosolic FGFR1 fusion mutants

**Supplementary Figure 4 Expression of all significant enriched pathways in cortical regions.** Expression is based on the mean expression of genes in the pathways. Gene expression was Z-scored across regions. Regions are sorted based on the *t*-statistics of ΔCT between Parkinson's disease patients and controls; positive scores are indicative of cortical atrophy. The pathways are sorted based on the average PLS regression coefficients of genes involved in the pathway. BH-corrected *P*-values for all significantly enriched pathways are in Supplementary Table 4. * = APC:Cdc20 mediated degradation of cell cycle proteins prior to satisfation of the cell cycle checkpoint. ** = APC/C:Cdh1 mediated degradation of Cdc20 and other APC/C:Cdh1 targeted proteins in late mitosis/early G1.

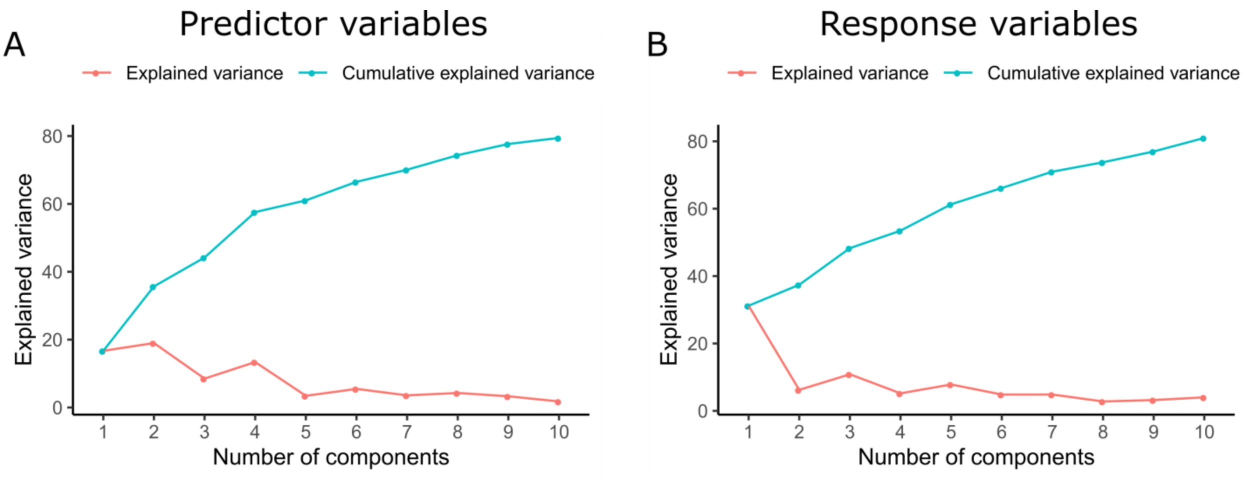


**Supplementary Figure 5 Explained variance in (A) predictor and (B) response variables of PLS *model-2*.**

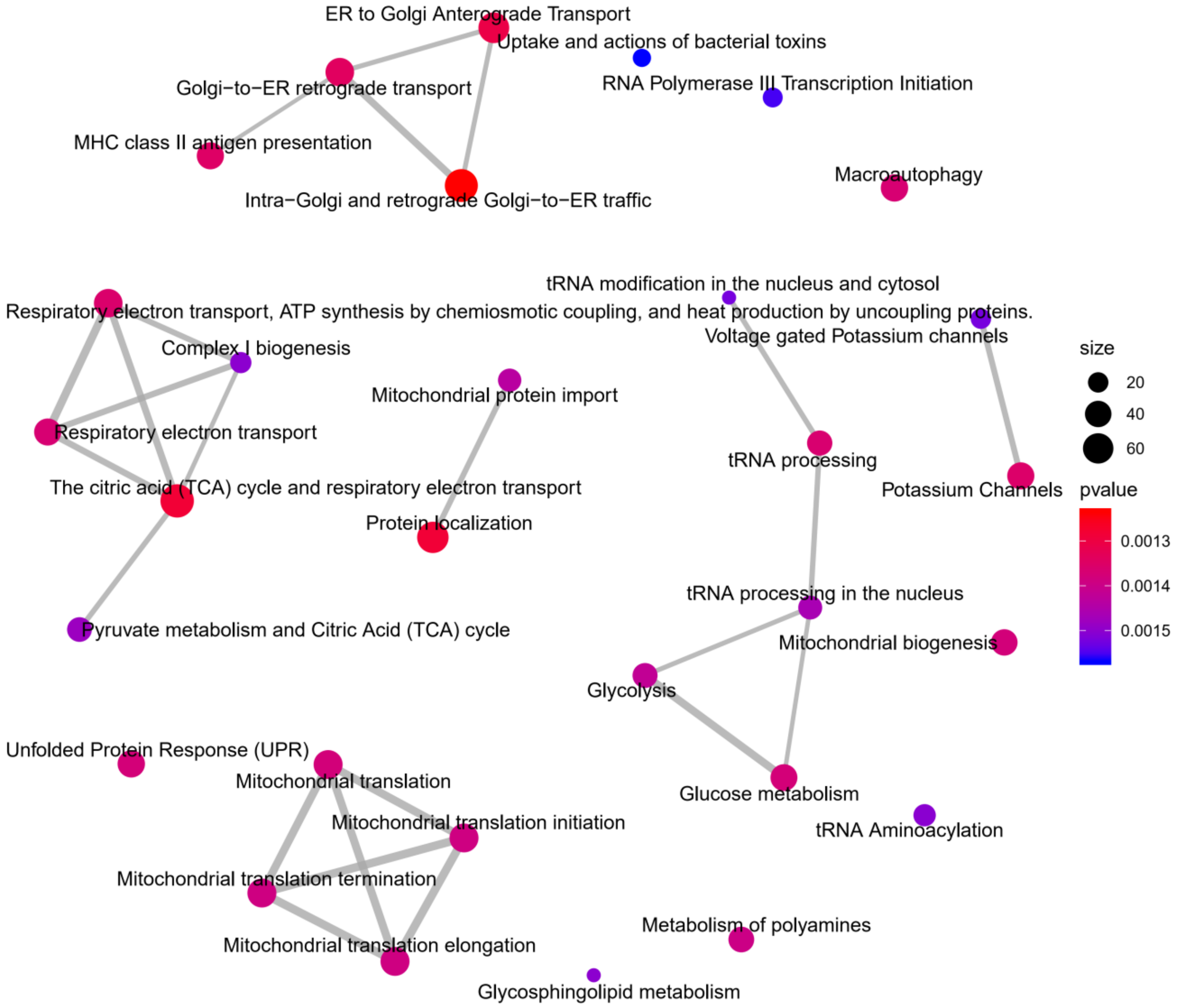


**Supplementary Figure 6 Top 30 significantly enriched pathways based on the PLS gene weights for *component-1* of *model-2* (BH-corrected $P < 0.05$).** Node size indicates the number of genes in the pathway, node color shows the unadjusted *P*-values, and edge thickness indicates the number of overlapping genes between two pathways. A list of all enriched pathways is given in Supplementary Table 5.

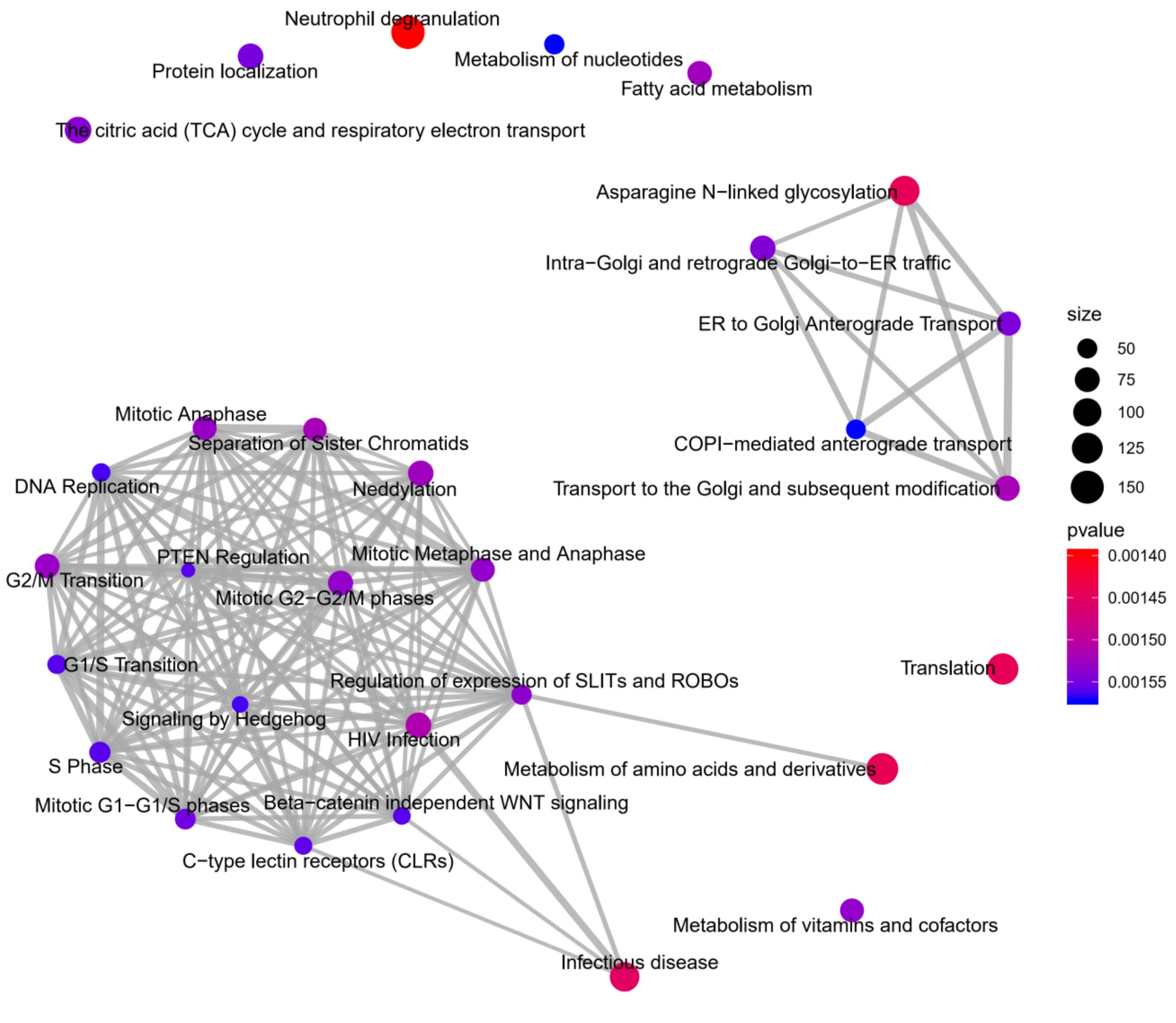


**Supplementary Figure 7 Top 30 significantly enriched pathways based on the PLS gene weights for *component-2* of *model-2* (BH-corrected *P* < 0.05).** Node size indicates the number of genes in the pathway, node color shows the unadjusted *P*-values, and edge thickness indicates the number of overlapping genes between two pathways. A list of all enriched pathways is given in Supplementary Table 6.

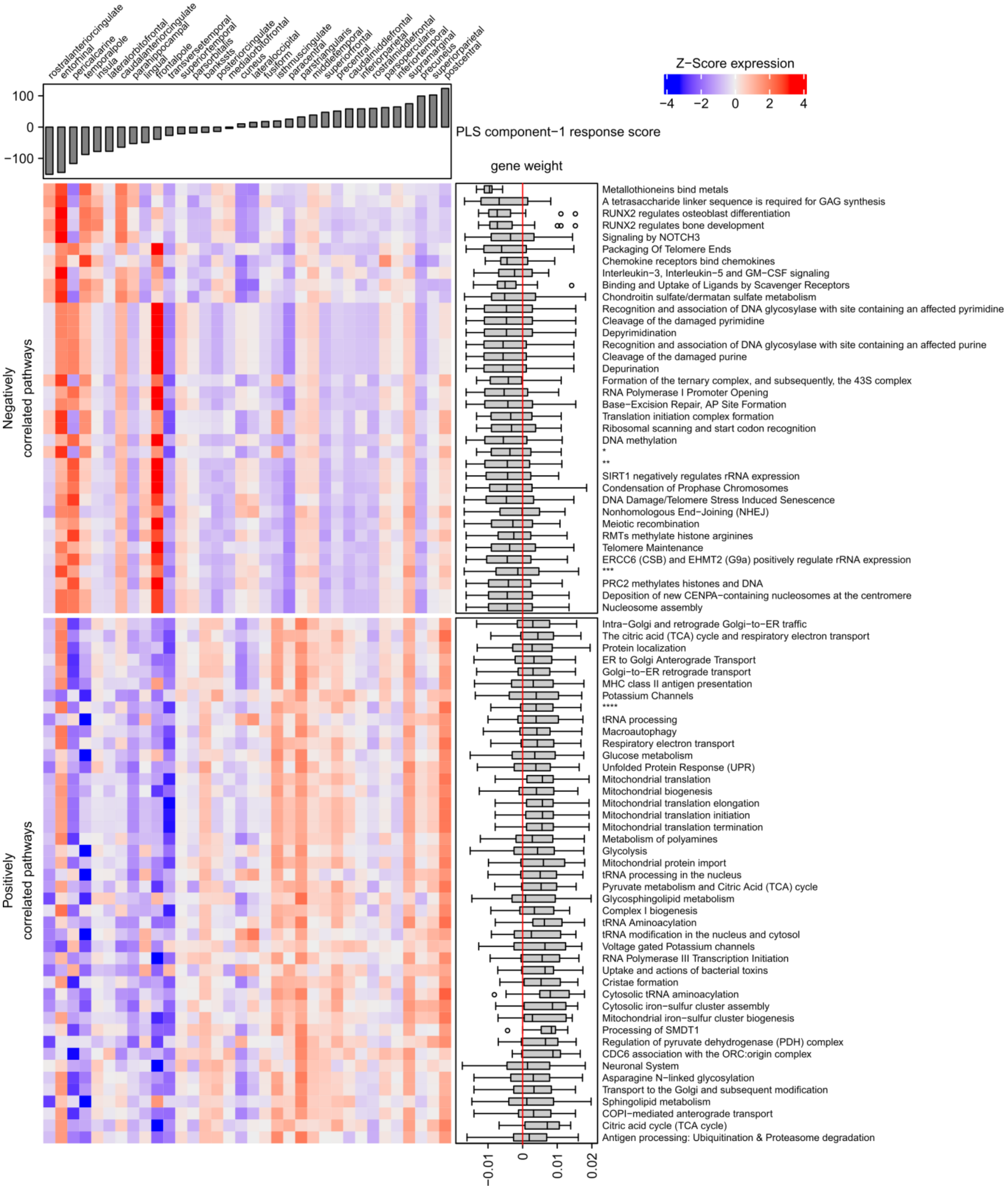

rostralanteriorcingulate
entorhinal
pericalcarine
temporalpole
insula
lateralorbitofrontal
caudalanteriorcingulate
parahippocampal
lingual
frontalpole
transversetemporal
superiortemporal
parsorbitalis
bankssts
posteriorcingulate
medialorbitofrontal
cuneus
lateraloccipital
fusiform
isthmuscingulate
paracentral
parstriangularis
middletemporal
superiorfrontal
precentral
caudalmiddlefrontal
inferiorparietal
rostralmiddlefrontal
parsopercularis
inferiortemporal
supramarginal
precuneus
superiorparietal
postcentral
Z-Score expression
−4
−2
0
2
4
100
0
−100
PLS component−1 response score
gene weight
Negatively correlated pathways
Metallothioneins bind metals
A tetrasaccharide linker sequence is required for GAG synthesis
RUNX2 regulates osteoblast differentiation
RUNX2 regulates bone development
Signaling by NOTCH3
Packaging Of Telomere Ends
Chemokine receptors bind chemokines
Interleukin−3, Interleukin−5 and GM−CSF signaling
Binding and Uptake of Ligands by Scavenger Receptors
Chondroitin sulfate/dermatan sulfate metabolism
Recognition and association of DNA glycosylase with site containing an affected pyrimidine
Cleavage of the damaged pyrimidine
Depyrimidination
Recognition and association of DNA glycosylase with site containing an affected purine
Cleavage of the damaged purine
Depurination
Formation of the ternary complex, and subsequently, the 43S complex
RNA Polymerase I Promoter Opening
Base−Excision Repair, AP Site Formation
Translation initiation complex formation
Ribosomal scanning and start codon recognition
DNA methylation
*
**
SIRT1 negatively regulates rRNA expression
Condensation of Prophase Chromosomes
DNA Damage/Telomere Stress Induced Senescence
Nonhomologous End−Joining (NHEJ)
Meiotic recombination
RMTs methylate histone arginines
Telomere Maintenance
ERCC6 (CSB) and EHMT2 (G9a) positively regulate rRNA expression
***
PRC2 methylates histones and DNA
Deposition of new CENPA−containing nucleosomes at the centromere
Nucleosome assembly
Positively correlated pathways
Intra−Golgi and retrograde Golgi−to−ER traffic
The citric acid (TCA) cycle and respiratory electron transport
Protein localization
ER to Golgi Anterograde Transport
Golgi−to−ER retrograde transport
MHC class II antigen presentation
Potassium Channels
****
tRNA processing
Macroautophagy
Respiratory electron transport
Glucose metabolism
Unfolded Protein Response (UPR)
Mitochondrial translation
Mitochondrial biogenesis
Mitochondrial translation elongation
Mitochondrial translation initiation
Mitochondrial translation termination
Metabolism of polyamines
Glycolysis
Mitochondrial protein import
tRNA processing in the nucleus
Pyruvate metabolism and Citric Acid (TCA) cycle
Glycosphingolipid metabolism
Complex I biogenesis
tRNA Aminoacylation
tRNA modification in the nucleus and cytosol
Voltage gated Potassium channels
RNA Polymerase III Transcription Initiation
Uptake and actions of bacterial toxins
Cristae formation
Cytosolic tRNA aminoacylation
Cytosolic iron−sulfur cluster assembly
Mitochondrial iron−sulfur cluster biogenesis
Processing of SMDT1
Regulation of pyruvate dehydrogenase (PDH) complex
CDC6 association with the ORC:origin complex
Neuronal System
Asparagine N−linked glycosylation
Transport to the Golgi and subsequent modification
Sphingolipid metabolism
COPI−mediated anterograde transport
Citric acid cycle (TCA cycle)
Antigen processing: Ubiquitination & Proteasome degradation
−0.01
0
0.01
0.02

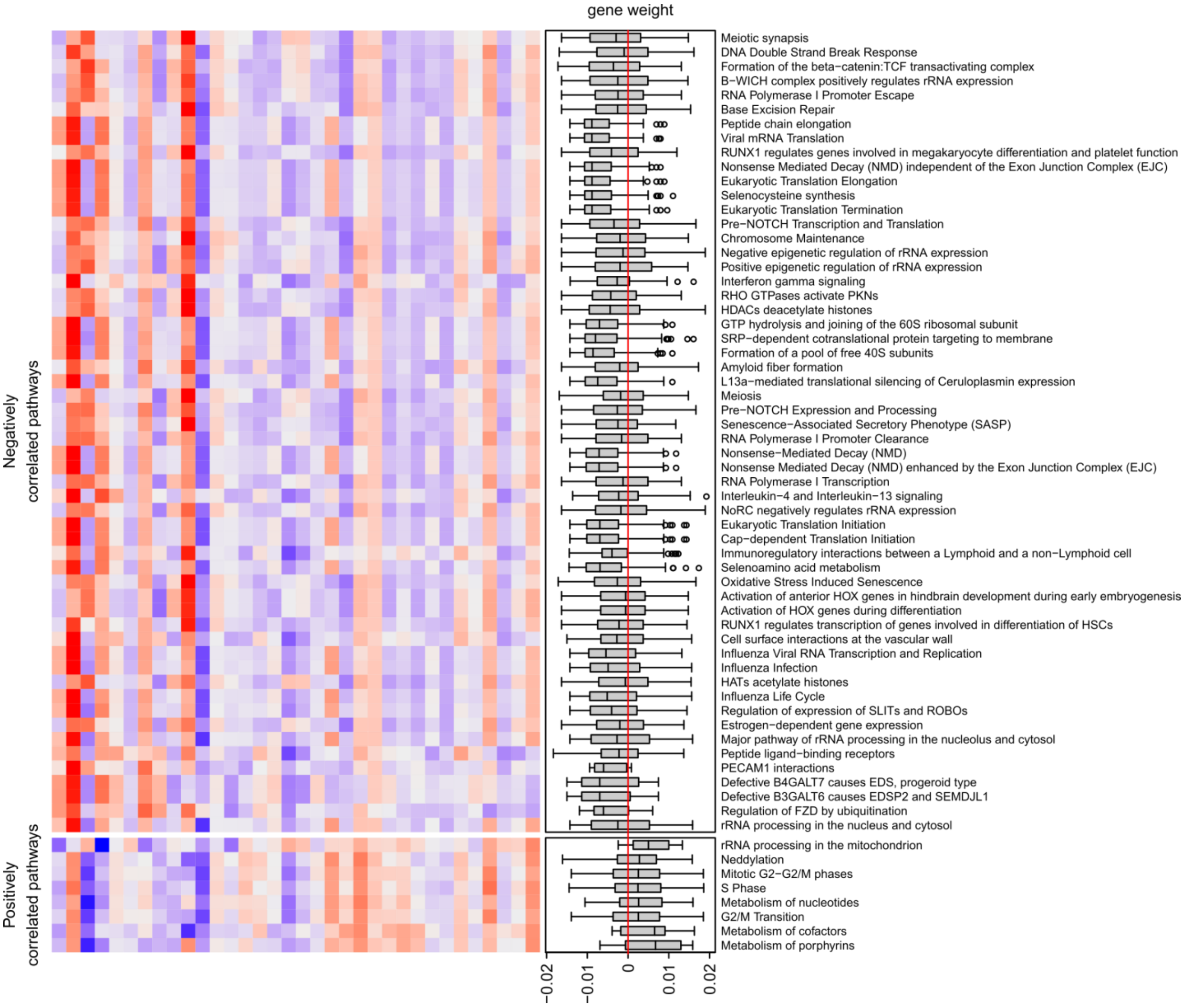


**Supplementary Figure 8 Expression of all significant enriched pathways in cortical regions in PLS *model-2* for *component-1*.** Expression is based on the mean expression of genes in the pathways. Gene expression was Z-scored across regions. Regions are sorted based on the *t*-statistics of ΔCT between Parkinson's disease patients and controls; positive scores are indicative of cortical atrophy. The pathways are sorted based on the average PLS regression coefficients of genes involved in the pathway. BH-corrected *P*-values for all significantly enriched pathways are in Supplementary Table X. * = Activation of the mRNA upon binding of the cap−binding complex and eIFs, and subsequent binding to 43S, ** = Activated PKN1 stimulates transcription of AR (androgen receptor) regulated genes KLK2 and KLK3. *** = Recruitment and ATM−mediated phosphorylation of repair and signaling proteins at DNA double strand breaks. **** = Respiratory electron transport, ATP synthesis by chemiosmotic coupling, and heat production by uncoupling proteins.

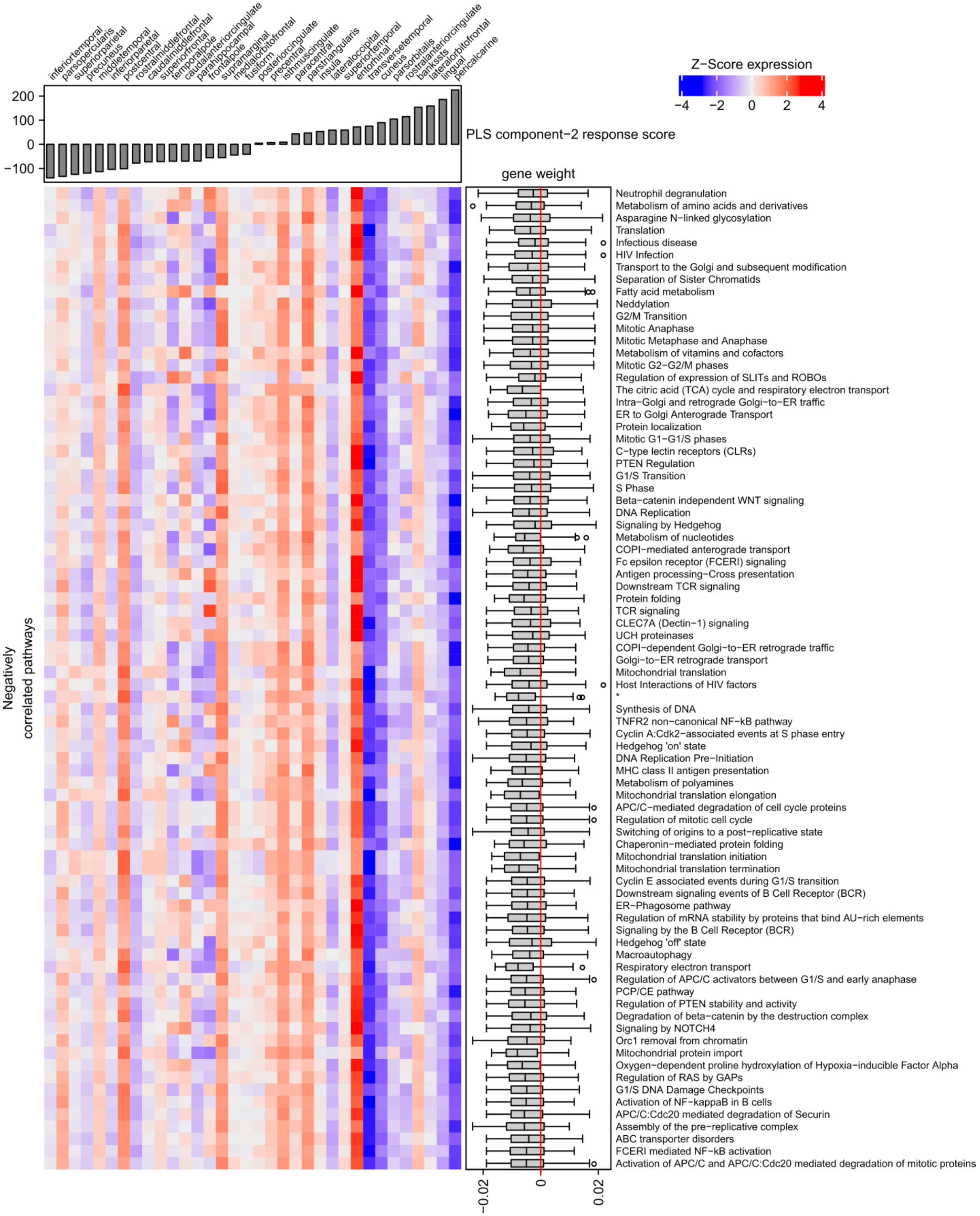
inferiortemporal
parsopercularis
superiorparietal
precuneus
middletemporal
inferiorparietal
postcentral
rostralmiddlefrontal
caudalmiddlefrontal
superiorfrontal
temporalpole
caudalanteriorcingulate
parahippocampal
frontalpole
supramarginal
medialorbitofrontal
fusiform
posteriorcingulate
precentral
isthmuscingulate
paracentral
parstriangularis
insula
lateraloccipital
superiortemporal
entorhinal
transversetemporal
cuneus
parsorbitalis
rostralanteriorcingulate
bankssts
lateralorbitofrontal
lingual
pericalcarine
200
100
0
−100
PLS component−2 response score
Z−Score expression
−4
−2
0
2
4
gene weight
Negatively correlated pathways
Neutrophil degranulation
Metabolism of amino acids and derivatives
Asparagine N−linked glycosylation
Translation
Infectious disease
HIV Infection
Transport to the Golgi and subsequent modification
Separation of Sister Chromatids
Fatty acid metabolism
Neddylation
G2/M Transition
Mitotic Anaphase
Mitotic Metaphase and Anaphase
Metabolism of vitamins and cofactors
Mitotic G2−G2/M phases
Regulation of expression of SLITs and ROBOs
The citric acid (TCA) cycle and respiratory electron transport
Intra−Golgi and retrograde Golgi−to−ER traffic
ER to Golgi Anterograde Transport
Protein localization
Mitotic G1−G1/S phases
C−type lectin receptors (CLRs)
PTEN Regulation
G1/S Transition
S Phase
Beta−catenin independent WNT signaling
DNA Replication
Signaling by Hedgehog
Metabolism of nucleotides
COPI−mediated anterograde transport
Fc epsilon receptor (FCERI) signaling
Antigen processing−Cross presentation
Downstream TCR signaling
Protein folding
TCR signaling
CLEC7A (Dectin−1) signaling
UCH proteinases
COPI−dependent Golgi−to−ER retrograde traffic
Golgi−to−ER retrograde transport
Mitochondrial translation
Host Interactions of HIV factors
*
Synthesis of DNA
TNFR2 non−canonical NF−kB pathway
Cyclin A:Cdk2−associated events at S phase entry
Hedgehog 'on' state
DNA Replication Pre−Initiation
MHC class II antigen presentation
Metabolism of polyamines
Mitochondrial translation elongation
APC/C−mediated degradation of cell cycle proteins
Regulation of mitotic cell cycle
Switching of origins to a post−replicative state
Chaperonin−mediated protein folding
Mitochondrial translation initiation
Mitochondrial translation termination
Cyclin E associated events during G1/S transition
Downstream signaling events of B Cell Receptor (BCR)
ER−Phagosome pathway
Regulation of mRNA stability by proteins that bind AU−rich elements
Signaling by the B Cell Receptor (BCR)
Hedgehog 'off' state
Macroautophagy
Respiratory electron transport
Regulation of APC/C activators between G1/S and early anaphase
PCP/CE pathway
Regulation of PTEN stability and activity
Degradation of beta−catenin by the destruction complex
Signaling by NOTCH4
Orc1 removal from chromatin
Mitochondrial protein import
Oxygen−dependent proline hydroxylation of Hypoxia−inducible Factor Alpha
Regulation of RAS by GAPs
G1/S DNA Damage Checkpoints
Activation of NF−kappaB in B cells
APC/C:Cdc20 mediated degradation of Securin
Assembly of the pre−replicative complex
ABC transporter disorders
FCERI mediated NF−kB activation
Activation of APC/C and APC/C:Cdc20 mediated degradation of mitotic proteins
−0.02
0
0.02

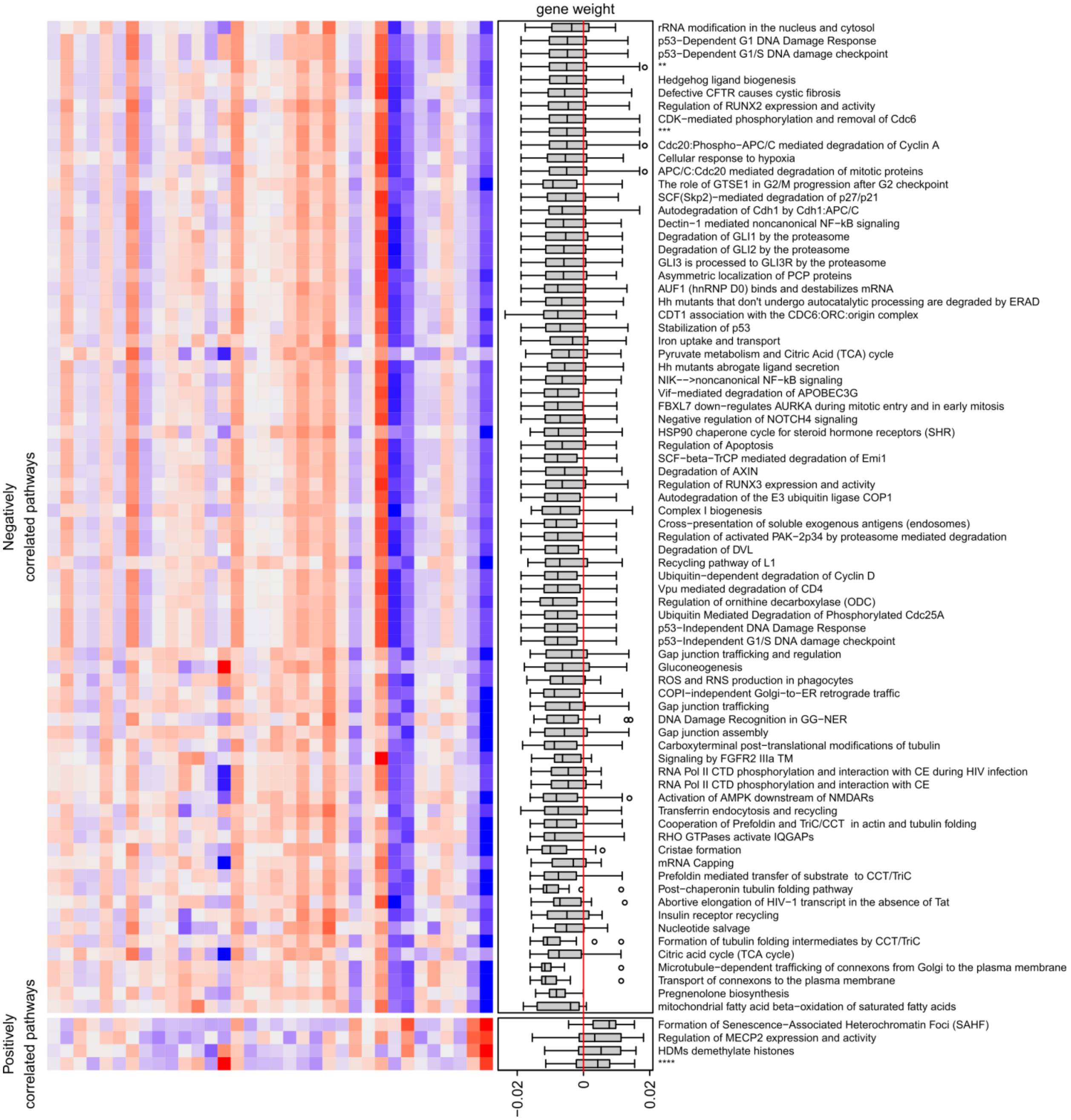

gene weight
rRNA modification in the nucleus and cytosol
p53−Dependent G1 DNA Damage Response
p53−Dependent G1/S DNA damage checkpoint
**
Hedgehog ligand biogenesis
Defective CFTR causes cystic fibrosis
Regulation of RUNX2 expression and activity
CDK−mediated phosphorylation and removal of Cdc6
***
Cdc20:Phospho−APC/C mediated degradation of Cyclin A
Cellular response to hypoxia
APC/C:Cdc20 mediated degradation of mitotic proteins
The role of GTSE1 in G2/M progression after G2 checkpoint
SCF(Skp2)−mediated degradation of p27/p21
Autodegradation of Cdh1 by Cdh1:APC/C
Dectin−1 mediated noncanonical NF−kB signaling
Degradation of GLI1 by the proteasome
Degradation of GLI2 by the proteasome
GLI3 is processed to GLI3R by the proteasome
Asymmetric localization of PCP proteins
AUF1 (hnRNP D0) binds and destabilizes mRNA
Hh mutants that don't undergo autocatalytic processing are degraded by ERAD
CDT1 association with the CDC6:ORC:origin complex
Stabilization of p53
Iron uptake and transport
Pyruvate metabolism and Citric Acid (TCA) cycle
Hh mutants abrogate ligand secretion
NIK−−>noncanonical NF−kB signaling
Vif−mediated degradation of APOBEC3G
FBXL7 down−regulates AURKA during mitotic entry and in early mitosis
Negative regulation of NOTCH4 signaling
HSP90 chaperone cycle for steroid hormone receptors (SHR)
Regulation of Apoptosis
SCF−beta−TrCP mediated degradation of Emi1
Degradation of AXIN
Regulation of RUNX3 expression and activity
Autodegradation of the E3 ubiquitin ligase COP1
Complex I biogenesis
Cross−presentation of soluble exogenous antigens (endosomes)
Regulation of activated PAK−2p34 by proteasome mediated degradation
Degradation of DVL
Recycling pathway of L1
Ubiquitin−dependent degradation of Cyclin D
Vpu mediated degradation of CD4
Regulation of ornithine decarboxylase (ODC)
Ubiquitin Mediated Degradation of Phosphorylated Cdc25A
p53−Independent DNA Damage Response
p53−Independent G1/S DNA damage checkpoint
Gap junction trafficking and regulation
Gluconeogenesis
ROS and RNS production in phagocytes
COPI−independent Golgi−to−ER retrograde traffic
Gap junction trafficking
DNA Damage Recognition in GG−NER
Gap junction assembly
Carboxyterminal post−translational modifications of tubulin
Signaling by FGFR2 IIIa TM
RNA Pol II CTD phosphorylation and interaction with CE during HIV infection
RNA Pol II CTD phosphorylation and interaction with CE
Activation of AMPK downstream of NMDARs
Transferrin endocytosis and recycling
Cooperation of Prefoldin and TriC/CCT in actin and tubulin folding
RHO GTPases activate IQGAPs
Cristae formation
mRNA Capping
Prefoldin mediated transfer of substrate to CCT/TriC
Post−chaperonin tubulin folding pathway
Abortive elongation of HIV−1 transcript in the absence of Tat
Insulin receptor recycling
Nucleotide salvage
Formation of tubulin folding intermediates by CCT/TriC
Citric acid cycle (TCA cycle)
Microtubule−dependent trafficking of connexons from Golgi to the plasma membrane
Transport of connexons to the plasma membrane
Pregnenolone biosynthesis
mitochondrial fatty acid beta−oxidation of saturated fatty acids
Formation of Senescence−Associated Heterochromatin Foci (SAHF)
Regulation of MECP2 expression and activity
HDMs demethylate histones
****
Negatively correlated pathways
Positively correlated pathways
−0.02
0
0.02

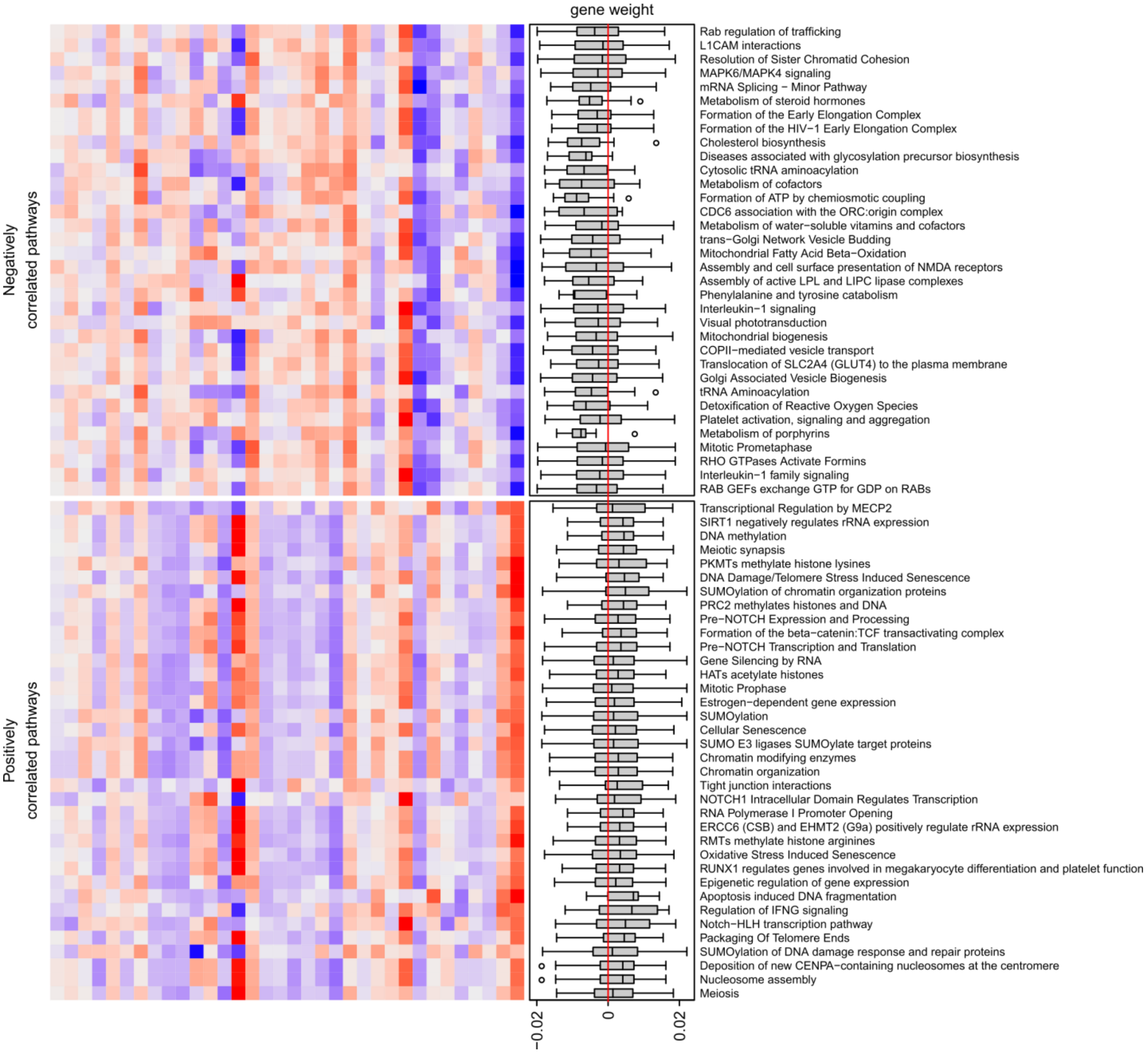


**Supplementary Figure 9 Expression of all significant enriched pathways in cortical regions in PLS *model-2* for *component-2*.** Expression is based on the mean expression of genes in the pathways. Gene expression was Z-scored across regions. Regions are sorted based on the *t*-statistics of ΔCT between Parkinson's disease patients and controls; positive scores are indicative of cortical atrophy. The pathways are sorted based on the average PLS regression coefficients of genes involved in the pathway. BH-corrected *P*-values for all significantly enriched pathways are in Supplementary Table X. * = Respiratory electron transport, ATP synthesis by chemiosmotic coupling, and heat production by uncoupling proteins. ** = APC:Cdc20 mediated degradation of cell cycle proteins prior to satisfation of the cell cycle checkpoint. *** = APC/C:Cdh1 mediated degradation of Cdc20 and other APC/C:Cdh1 targeted proteins in late mitosis/early G1. **** = Activated PKN1 stimulates transcription of AR (androgen receptor) regulated genes KLK2 and KLK3.

### Supplementary Tables

**Supplementary Table 1 All 68 cortical regions from both hemispheres with cortical thickness differences (ΔCT) in age-matched controls compared to Parkinson's disease patients.** Regions are sorted based on the BH-corrected *P* and "lh" and "rh" indicates the left or right hemisphere, respectively. PD = Parkinson's disease.

| Region ID | Region name | *t*-score | ΔCT | Mean CT control | Mean CT PD | BH-corrected P |
|---|---|---|---|---|---|---|
| 10 | lh_lateraloccipital | 4.29 | 0.05 | 2.03 | 1.97 | 1.66e-03 |
| 58 | rh_parsopercularis | -3.98 | -0.07 | 2.41 | 2.48 | 2.98e-03 |
| 51 | rh_lateraloccipital | 3.70 | 0.05 | 2.07 | 2.03 | 5.88e-03 |
| 17 | lh_parsopercularis | -3.26 | -0.05 | 2.42 | 2.48 | 2.00e-02 |
| 67 | rh_rostralmiddlefrontal | -3.21 | -0.05 | 2.26 | 2.31 | 2.00e-02 |
| 2 | lh_caudalanteriorcingulate | 3.12 | 0.09 | 2.84 | 2.75 | 2.25e-02 |
| 73 | rh_temporalpole | -3.07 | -0.10 | 3.41 | 3.51 | 2.25e-02 |
| 70 | rh_superiortemporal | -2.96 | -0.05 | 2.60 | 2.65 | 2.84e-02 |
| 50 | rh_isthmuscingulate | 2.89 | 0.06 | 2.29 | 2.24 | 3.14e-02 |
| 61 | rh_pericalcarine | 2.77 | 0.03 | 1.39 | 1.36 | 4.09e-02 |
| 48 | rh_inferiorparietal | 2.66 | 0.04 | 2.34 | 2.30 | 5.13e-02 |
| 53 | rh_lingual | 2.61 | 0.03 | 1.80 | 1.77 | 5.53e-02 |
| 7 | lh_inferiorparietal | 2.54 | 0.04 | 2.31 | 2.27 | 5.53e-02 |
| 15 | lh_parahippocampal | 2.52 | 0.08 | 2.51 | 2.42 | 5.53e-02 |
| 26 | lh_rostralmiddlefrontal | -2.53 | -0.04 | 2.27 | 2.31 | 5.53e-02 |
| 5 | lh_entorhinal | -2.47 | -0.09 | 3.11 | 3.20 | 5.85e-02 |
| 9 | lh_isthmuscingulate | 2.46 | 0.05 | 2.32 | 2.28 | 5.85e-02 |
| 46 | rh_entorhinal | -2.32 | -0.08 | 3.27 | 3.36 | 7.87e-02 |
| 19 | lh_parstriangularis | -2.27 | -0.04 | 2.30 | 2.34 | 8.12e-02 |
| 22 | lh_posteriorcingulate | 2.29 | 0.04 | 2.45 | 2.40 | 8.12e-02 |
| 56 | rh_parahippocampal | 1.94 | 0.06 | 2.48 | 2.43 | 1.74e-01 |
| 62 | rh_postcentral | -1.91 | -0.03 | 1.88 | 1.91 | 1.76e-01 |
| 32 | lh_temporalpole | -1.81 | -0.06 | 3.32 | 3.38 | 2.10e-01 |
| 64 | rh_precentral | -1.79 | -0.03 | 2.37 | 2.40 | 2.10e-01 |
| 63 | rh_posteriorcingulate | 1.53 | 0.03 | 2.45 | 2.42 | 3.47e-01 |
| 20 | lh_pericalcarine | 1.49 | 0.01 | 1.36 | 1.35 | 3.51e-01 |
| 21 | lh_postcentral | -1.48 | -0.02 | 1.92 | 1.95 | 3.51e-01 |
| 29 | lh_superiortemporal | -1.35 | -0.02 | 2.59 | 2.61 | 4.32e-01 |
| 1 | lh_bankssts | -1.31 | -0.02 | 2.30 | 2.32 | 4.33e-01 |
| 44 | rh_caudalmiddlefrontal | -1.31 | -0.02 | 2.43 | 2.46 | 4.33e-01 |
| 30 | lh_supramarginal | 1.25 | 0.02 | 2.43 | 2.41 | 4.53e-01 |
| 74 | rh_transversetemporal | 1.25 | 0.03 | 2.15 | 2.12 | 4.53e-01 |
| 42 | rh_bankssts | -1.21 | -0.02 | 2.40 | 2.42 | 4.57e-01 |
| 57 | rh_paracentral | 1.22 | 0.03 | 2.23 | 2.20 | 4.57e-01 |
| 3 | lh_caudalmiddlefrontal | -1.06 | -0.02 | 2.46 | 2.48 | 5.37e-01 |
| 11 | lh_lateralorbitofrontal | -1.06 | -0.02 | 2.56 | 2.58 | 5.37e-01 |
| 75 | rh_insula | -1.08 | -0.02 | 2.86 | 2.88 | 5.37e-01 |
| 25 | lh_rostralanteriorcingulate | 0.98 | 0.02 | 2.87 | 2.84 | 5.62e-01 |
| 43 | rh_caudalanteriorcingulate | 0.97 | 0.03 | 2.69 | 2.67 | 5.62e-01 |
| 52 | rh_lateralorbitofrontal | 1.00 | 0.02 | 2.53 | 2.51 | 5.62e-01 |
| 71 | rh_supramarginal | 0.92 | 0.02 | 2.43 | 2.41 | 5.92e-01 |
| 23 | lh_precentral | -0.88 | -0.02 | 2.40 | 2.42 | 6.12e-01 |
| 13 | lh_medialorbitofrontal | -0.83 | -0.01 | 2.38 | 2.39 | 6.40e-01 |
| 12 | lh_lingual | 0.78 | 0.01 | 1.75 | 1.74 | 6.47e-01 |
| 34 | lh_insula | -0.78 | -0.02 | 2.89 | 2.91 | 6.47e-01 |
| 59 | rh_parsorbitalis | 0.80 | 0.02 | 2.57 | 2.55 | 6.47e-01 |
| 18 | lh_parsorbitalis | -0.71 | -0.02 | 2.54 | 2.56 | 6.88e-01 |
| 72 | rh_frontalpole | -0.48 | -0.02 | 2.68 | 2.70 | 8.94e-01 |
| 65 | rh_precuneus | 0.46 | 0.01 | 2.19 | 2.18 | 9.01e-01 |
| 14 | lh_middletemporal | 0.43 | 0.01 | 2.75 | 2.74 | 9.06e-01 |
| 28 | lh_superiorparietal | -0.41 | -0.01 | 2.01 | 2.02 | 9.06e-01 |

| | | | | | | |
|---|---|---|---|---|---|---|
| 27 | lh_superiorfrontal | -0.36 | -0.01 | 2.66 | 2.67 | 9.27e-01 |
| 66 | rh_rostralanteriorcingulate | 0.36 | 0.01 | 2.87 | 2.86 | 9.27e-01 |
| 4 | lh_cuneus | 0.28 | 0.00 | 1.61 | 1.61 | 9.30e-01 |
| 6 | lh_fusiform | -0.26 | 0.00 | 2.47 | 2.48 | 9.30e-01 |
| 49 | rh_inferiortemporal | -0.27 | 0.00 | 2.65 | 2.66 | 9.30e-01 |
| 54 | rh_medialorbitofrontal | 0.33 | 0.01 | 2.47 | 2.46 | 9.30e-01 |
| 69 | rh_superiorparietal | -0.30 | 0.00 | 1.99 | 2.00 | 9.30e-01 |
| 55 | rh_middletemporal | -0.20 | 0.00 | 2.76 | 2.76 | 9.69e-01 |
| 8 | lh_inferiortemporal | -0.17 | 0.00 | 2.66 | 2.66 | 9.70e-01 |
| 16 | lh_paracentral | -0.14 | 0.00 | 2.21 | 2.21 | 9.70e-01 |
| 24 | lh_precuneus | 0.06 | 0.00 | 2.16 | 2.16 | 9.70e-01 |
| 31 | lh_frontalpole | -0.08 | 0.00 | 2.74 | 2.74 | 9.70e-01 |
| 45 | rh_cuneus | 0.06 | 0.00 | 1.62 | 1.62 | 9.70e-01 |
| 47 | rh_fusiform | 0.08 | 0.00 | 2.47 | 2.46 | 9.70e-01 |
| 60 | rh_parstriangularis | -0.10 | 0.00 | 2.32 | 2.33 | 9.70e-01 |
| 68 | rh_superiorfrontal | -0.06 | 0.00 | 2.66 | 2.66 | 9.70e-01 |
| 33 | lh_transversetemporal | -0.04 | 0.00 | 2.09 | 2.09 | 9.71e-01 |

**Supplementary Table 2 Cortical thickness differences (ΔCT) of the left hemisphere compared to the right in Parkinson's disease patients.** Regions are sorted based on the BH-corrected *P*.

| Region ID | Region name | *t*-score | ΔCT | Mean CT left hemisphere | Mean CT right hemisphere | BH-corrected P |
|---|---|---|---|---|---|---|
| 1 | lh_bankssts | 4.52 | 0.10 | 2.42 | 2.32 | 3.02e-04 |
| 5 | lh_entorhinal | 3.60 | 0.15 | 3.36 | 3.20 | 1.25e-02 |
| 32 | lh_temporalpole | 3.43 | 0.12 | 3.51 | 3.38 | 2.24e-02 |
| 13 | lh_medialorbitofrontal | 3.40 | 0.07 | 2.46 | 2.39 | 2.35e-02 |
| 10 | lh_lateraloccipital | 3.31 | 0.05 | 2.03 | 1.97 | 3.10e-02 |
| 11 | lh_lateralorbitofrontal | -3.21 | -0.06 | 2.51 | 2.58 | 4.21e-02 |
| 2 | lh_caudalanteriorcingulate | -2.65 | -0.09 | 2.67 | 2.75 | 2.35e-01 |
| 3 | lh_caudalmiddlefrontal | -0.86 | -0.02 | 2.46 | 2.48 | 1.00e+00 |
| 4 | lh_cuneus | 0.68 | 0.01 | 1.62 | 1.61 | 1.00e+00 |
| 6 | lh_fusiform | -0.79 | -0.02 | 2.46 | 2.48 | 1.00e+00 |
| 7 | lh_inferiorparietal | 1.42 | 0.03 | 2.30 | 2.27 | 1.00e+00 |
| 8 | lh_inferiortemporal | -0.09 | 0.00 | 2.66 | 2.66 | 1.00e+00 |
| 9 | lh_isthmuscingulate | -1.80 | -0.04 | 2.24 | 2.28 | 1.00e+00 |
| 12 | lh_lingual | 1.49 | 0.02 | 1.77 | 1.74 | 1.00e+00 |
| 14 | lh_middletemporal | 1.13 | 0.02 | 2.76 | 2.74 | 1.00e+00 |
| 15 | lh_parahippocampal | 0.03 | 0.00 | 2.43 | 2.42 | 1.00e+00 |
| 16 | lh_paracentral | -0.28 | -0.01 | 2.20 | 2.21 | 1.00e+00 |
| 17 | lh_parsopercularis | 0.31 | 0.01 | 2.48 | 2.48 | 1.00e+00 |
| 18 | lh_parsorbitalis | -0.39 | -0.01 | 2.55 | 2.56 | 1.00e+00 |
| 19 | lh_parstriangularis | -0.86 | -0.02 | 2.33 | 2.34 | 1.00e+00 |
| 20 | lh_pericalcarine | 1.17 | 0.01 | 1.36 | 1.35 | 1.00e+00 |
| 21 | lh_postcentral | -1.79 | -0.03 | 1.91 | 1.95 | 1.00e+00 |
| 22 | lh_posteriorcingulate | 0.88 | 0.02 | 2.42 | 2.40 | 1.00e+00 |
| 23 | lh_precentral | -0.76 | -0.02 | 2.40 | 2.42 | 1.00e+00 |
| 24 | lh_precuneus | 0.86 | 0.02 | 2.18 | 2.16 | 1.00e+00 |
| 25 | lh_rostralanteriorcingulate | 0.65 | 0.02 | 2.86 | 2.84 | 1.00e+00 |
| 26 | lh_rostralmiddlefrontal | -0.01 | 0.00 | 2.31 | 2.31 | 1.00e+00 |
| 27 | lh_superiorfrontal | -0.42 | -0.01 | 2.66 | 2.67 | 1.00e+00 |
| 28 | lh_superiorparietal | -1.01 | -0.02 | 2.00 | 2.02 | 1.00e+00 |
| 29 | lh_superiortemporal | 1.93 | 0.04 | 2.65 | 2.61 | 1.00e+00 |
| 30 | lh_supramarginal | 0.27 | 0.01 | 2.41 | 2.41 | 1.00e+00 |
| 31 | lh_frontalpole | -1.04 | -0.04 | 2.70 | 2.74 | 1.00e+00 |
| 33 | lh_transversetemporal | 0.92 | 0.03 | 2.12 | 2.09 | 1.00e+00 |
| 34 | lh_insula | -1.21 | -0.03 | 2.88 | 2.91 | 1.00e+00 |

**Supplementary Table 3 Number of AHBA samples in ROIs from FreeSurfer.** All samples and segmented cortical regions are from the left hemisphere only.

| Region ID | Name | Donor 861 | Donor 10021 | Donor 12876 | Donor 14380 | Donor 15496 | Donor 15697 | Total |
|---|---|---|---|---|---|---|---|---|
| 1 | bankssts | 6 | 3 | 4 | 3 | 3 | 3 | 22 |
| 2 | caudalanteriorcingulate | 1 | 0 | 7 | 5 | 6 | 4 | 23 |
| 3 | caudalmiddlefrontal | 8 | 6 | 3 | 4 | 4 | 5 | 30 |
| 4 | cuneus | 2 | 2 | 2 | 6 | 5 | 4 | 21 |
| 5 | entorhinal | 5 | 2 | 3 | 5 | 2 | 8 | 25 |
| 6 | fusiform | 10 | 10 | 8 | 18 | 16 | 16 | 78 |
| 7 | inferiorparietal | 7 | 11 | 8 | 6 | 6 | 10 | 48 |
| 8 | inferiortemporal | 19 | 5 | 13 | 17 | 14 | 11 | 79 |
| 9 | isthmuscingulate | 0 | 4 | 3 | 5 | 6 | 3 | 21 |
| 10 | lateraloccipital | 3 | 9 | 0 | 12 | 8 | 10 | 42 |
| 11 | lateralorbitofrontal | 6 | 4 | 5 | 7 | 6 | 8 | 36 |
| 12 | lingual | 5 | 6 | 9 | 10 | 7 | 6 | 43 |
| 13 | medialorbitofrontal | 3 | 7 | 9 | 10 | 10 | 6 | 45 |
| 14 | middletemporal | 15 | 6 | 6 | 12 | 9 | 9 | 57 |
| 15 | parahippocampal | 7 | 2 | 3 | 5 | 5 | 3 | 25 |
| 16 | paracentral | 3 | 6 | 2 | 3 | 4 | 5 | 23 |
| 17 | parsopercularis | 5 | 1 | 0 | 3 | 6 | 5 | 20 |
| 18 | parsorbitalis | 0 | 0 | 2 | 6 | 6 | 2 | 16 |
| 19 | parstriangularis | 5 | 5 | 2 | 5 | 2 | 2 | 21 |
| 20 | pericalcarine | 1 | 2 | 1 | 0 | 5 | 6 | 15 |
| 21 | postcentral | 10 | 8 | 8 | 10 | 6 | 7 | 49 |
| 22 | posteriorcingulate | 7 | 6 | 6 | 10 | 4 | 7 | 40 |
| 23 | precentral | 16 | 9 | 10 | 12 | 8 | 10 | 65 |
| 24 | precuneus | 5 | 3 | 11 | 9 | 11 | 7 | 46 |
| 25 | rostralanteriorcingulate | 1 | 6 | 3 | 4 | 1 | 2 | 17 |
| 26 | rostralmiddlefrontal | 7 | 10 | 6 | 9 | 10 | 10 | 52 |
| 27 | superiorfrontal | 23 | 14 | 11 | 11 | 17 | 16 | 92 |
| 28 | superiorparietal | 8 | 8 | 11 | 13 | 9 | 12 | 61 |
| 29 | superiortemporal | 13 | 5 | 12 | 18 | 14 | 18 | 80 |
| 30 | supramarginal | 6 | 3 | 3 | 10 | 5 | 5 | 32 |
| 31 | frontalpole | 0 | 5 | 0 | 0 | 0 | 0 | 5 |
| 32 | temporalpole | 2 | 0 | 1 | 3 | 0 | 4 | 10 |
| 33 | transversetemporal | 2 | 2 | 1 | 1 | 1 | 0 | 7 |
| 34 | insula | 9 | 5 | 5 | 7 | 6 | 6 | 38 |

**Supplementary Table 4 Pathway enrichment of genes ranked by their weight for PLS *model-1* and *component-1*.**

| Pathway | P-value | BH-corrected *P*-value |
|---|---|---|
| Class C/3 (Metabotropic glutamate/pheromone receptors) | 1.91e-03 | 3.53e-02 |
| SUMOylation of chromatin organization proteins | 1.93e-03 | 3.53e-02 |
| Signaling by cytosolic FGFR1 fusion mutants | 2.01e-03 | 3.53e-02 |
| PCP/CE pathway | 2.02e-03 | 3.53e-02 |
| APC:Cdc20 mediated degradation of cell cycle proteins prior to satisfation of the cell cycle checkpoint | 2.03e-03 | 3.53e-02 |
| Hedgehog 'on' state | 2.03e-03 | 3.53e-02 |
| ABC transporter disorders | 2.05e-03 | 3.53e-02 |
| Cellular response to hypoxia | 2.05e-03 | 3.53e-02 |
| APC/C:Cdc20 mediated degradation of mitotic proteins | 2.05e-03 | 3.53e-02 |
| Activation of APC/C and APC/C:Cdc20 mediated degradation of mitotic proteins | 2.05e-03 | 3.53e-02 |
| Regulation of mRNA stability by proteins that bind AU-rich elements | 2.05e-03 | 3.53e-02 |
| Signaling by NOTCH4 | 2.05e-03 | 3.53e-02 |
| Downstream signaling events of B Cell Receptor (BCR) | 2.05e-03 | 3.53e-02 |
| ER-Phagosome pathway | 2.05e-03 | 3.53e-02 |
| Mitochondrial translation elongation | 2.06e-03 | 3.53e-02 |
| Metabolism of polyamines | 2.06e-03 | 3.53e-02 |
| Regulation of APC/C activators between G1/S and early anaphase | 2.06e-03 | 3.53e-02 |
| Switching of origins to a post-replicative state | 2.06e-03 | 3.53e-02 |

| | | |
|---|---|---|
| Mitochondrial translation initiation | 2.06e-03 | 3.53e-02 |
| Mitochondrial translation termination | 2.06e-03 | 3.53e-02 |
| APC/C-mediated degradation of cell cycle proteins | 2.06e-03 | 3.53e-02 |
| Regulation of mitotic cell cycle | 2.06e-03 | 3.53e-02 |
| CDK-mediated phosphorylation and removal of Cdc6 | 2.06e-03 | 3.53e-02 |
| APC/C:Cdh1 mediated degradation of Cdc20 and other APC/C:Cdh1 targeted proteins in late mitosis/early G1 | 2.06e-03 | 3.53e-02 |
| Cdc20:Phospho-APC/C mediated degradation of Cyclin A | 2.06e-03 | 3.53e-02 |
| Mitochondrial translation | 2.07e-03 | 3.53e-02 |
| Orc1 removal from chromatin | 2.07e-03 | 3.53e-02 |
| DNA Replication Pre-Initiation | 2.07e-03 | 3.53e-02 |
| Antigen processing-Cross presentation | 2.07e-03 | 3.53e-02 |
| Regulation of PTEN stability and activity | 2.08e-03 | 3.53e-02 |
| Ubiquitin-dependent degradation of Cyclin D | 2.08e-03 | 3.53e-02 |
| Vpu mediated degradation of CD4 | 2.08e-03 | 3.53e-02 |
| Regulation of ornithine decarboxylase (ODC) | 2.08e-03 | 3.53e-02 |
| CDT1 association with the CDC6:ORC:origin complex | 2.08e-03 | 3.53e-02 |
| Ubiquitin Mediated Degradation of Phosphorylated Cdc25A | 2.08e-03 | 3.53e-02 |
| p53-Independent DNA Damage Response | 2.08e-03 | 3.53e-02 |
| p53-Independent G1/S DNA damage checkpoint | 2.08e-03 | 3.53e-02 |
| The role of GTSE1 in G2/M progression after G2 checkpoint | 2.08e-03 | 3.53e-02 |
| Regulation of Apoptosis | 2.09e-03 | 3.53e-02 |
| Autodegradation of the E3 ubiquitin ligase COP1 | 2.09e-03 | 3.53e-02 |
| Regulation of RAS by GAPs | 2.10e-03 | 3.53e-02 |
| G1/S DNA Damage Checkpoints | 2.10e-03 | 3.53e-02 |
| Activation of NF-kappaB in B cells | 2.10e-03 | 3.53e-02 |
| APC/C:Cdc20 mediated degradation of Securin | 2.10e-03 | 3.53e-02 |
| Assembly of the pre-replicative complex | 2.10e-03 | 3.53e-02 |
| Vif-mediated degradation of APOBEC3G | 2.10e-03 | 3.53e-02 |
| FBXL7 down-regulates AURKA during mitotic entry and in early mitosis | 2.10e-03 | 3.53e-02 |
| Negative regulation of NOTCH4 signaling | 2.10e-03 | 3.53e-02 |
| Cross-presentation of soluble exogenous antigens (endosomes) | 2.11e-03 | 3.53e-02 |
| SCF-beta-TrCP mediated degradation of Emi1 | 2.11e-03 | 3.53e-02 |
| Regulation of activated PAK-2p34 by proteasome mediated degradation | 2.11e-03 | 3.53e-02 |
| AUF1 (hnRNP D0) binds and destabilizes mRNA | 2.11e-03 | 3.53e-02 |
| Degradation of AXIN | 2.11e-03 | 3.53e-02 |
| Hh mutants that don't undergo autocatalytic processing are degraded by ERAD | 2.11e-03 | 3.53e-02 |
| Stabilization of p53 | 2.11e-03 | 3.53e-02 |
| Regulation of RUNX3 expression and activity | 2.11e-03 | 3.53e-02 |
| Mitochondrial protein import | 2.12e-03 | 3.53e-02 |
| Oxygen-dependent proline hydroxylation of Hypoxia-inducible Factor Alpha | 2.12e-03 | 3.53e-02 |
| Hh mutants abrogate ligand secretion | 2.13e-03 | 3.53e-02 |
| NIK-->noncanonical NF-kB signaling | 2.13e-03 | 3.53e-02 |
| p53-Dependent G1 DNA Damage Response | 2.14e-03 | 3.53e-02 |
| p53-Dependent G1/S DNA damage checkpoint | 2.14e-03 | 3.53e-02 |
| Hedgehog ligand biogenesis | 2.14e-03 | 3.53e-02 |
| Mitotic Metaphase and Anaphase | 2.14e-03 | 3.53e-02 |
| Mitotic G2-G2/M phases | 2.14e-03 | 3.53e-02 |
| Defective CFTR causes cystic fibrosis | 2.14e-03 | 3.53e-02 |
| Peptide ligand-binding receptors | 2.15e-03 | 3.53e-02 |
| Degradation of DVL | 2.15e-03 | 3.53e-02 |
| Separation of Sister Chromatids | 2.17e-03 | 3.53e-02 |
| Regulation of expression of SLITs and ROBOs | 2.17e-03 | 3.53e-02 |
| DNA Damage Recognition in GG-NER | 2.18e-03 | 3.53e-02 |
| Mitotic Anaphase | 2.18e-03 | 3.53e-02 |
| G2/M Checkpoints | 2.18e-03 | 3.53e-02 |
| SCF(Skp2)-mediated degradation of p27/p21 | 2.18e-03 | 3.53e-02 |
| Autodegradation of Cdh1 by Cdh1:APC/C | 2.18e-03 | 3.53e-02 |
| Dectin-1 mediated noncanonical NF-kB signaling | 2.18e-03 | 3.53e-02 |
| Degradation of GLI1 by the proteasome | 2.18e-03 | 3.53e-02 |
| Degradation of GLI2 by the proteasome | 2.18e-03 | 3.53e-02 |
| GLI3 is processed to GLI3R by the proteasome | 2.18e-03 | 3.53e-02 |
| Protein localization | 2.18e-03 | 3.53e-02 |
| The citric acid (TCA) cycle and respiratory electron transport | 2.18e-03 | 3.53e-02 |
| Asymmetric localization of PCP proteins | 2.18e-03 | 3.53e-02 |
| Neddylation | 2.19e-03 | 3.53e-02 |
| Neutrophil degranulation | 2.20e-03 | 3.53e-02 |
| ER to Golgi Anterograde Transport | 2.22e-03 | 3.53e-02 |
| Asparagine N-linked glycosylation | 2.22e-03 | 3.53e-02 |
| Translation | 2.24e-03 | 3.53e-02 |

| | | |
|---|---|---|
| Signaling by Interleukins | 2.24e-03 | 3.53e-02 |
| Class A/1 (Rhodopsin-like receptors) | 2.28e-03 | 3.56e-02 |
| Metabolism of amino acids and derivatives | 2.33e-03 | 3.59e-02 |

**Supplementary Table 5 Pathway enrichment of genes ranked by their weight for PLS *model-2* and *component-1*.**

| **Pathway** | ***P*-value** | **BH-corrected *P*-value** |
|---|---|---|
| Intra-Golgi and retrograde Golgi-to-ER traffic | 1.24e-03 | 4.31e-02 |
| The citric acid (TCA) cycle and respiratory electron transport | 1.28e-03 | 4.31e-02 |
| Protein localization | 1.29e-03 | 4.31e-02 |
| ER to Golgi Anterograde Transport | 1.31e-03 | 4.31e-02 |
| Golgi-to-ER retrograde transport | 1.34e-03 | 4.31e-02 |
| MHC class II antigen presentation | 1.35e-03 | 4.31e-02 |
| Potassium Channels | 1.36e-03 | 4.31e-02 |
| Respiratory electron transport, ATP synthesis by chemiosmotic coupling, and heat production by uncoupling proteins. | 1.36e-03 | 4.31e-02 |
| tRNA processing | 1.36e-03 | 4.31e-02 |
| Macroautophagy | 1.37e-03 | 4.31e-02 |
| Respiratory electron transport | 1.37e-03 | 4.31e-02 |
| Glucose metabolism | 1.38e-03 | 4.31e-02 |
| Unfolded Protein Response (UPR) | 1.38e-03 | 4.31e-02 |
| Mitochondrial translation | 1.38e-03 | 4.31e-02 |
| Mitochondrial biogenesis | 1.38e-03 | 4.31e-02 |
| Mitochondrial translation elongation | 1.39e-03 | 4.31e-02 |
| Mitochondrial translation initiation | 1.39e-03 | 4.31e-02 |
| Mitochondrial translation termination | 1.39e-03 | 4.31e-02 |
| Metabolism of polyamines | 1.40e-03 | 4.31e-02 |
| Glycolysis | 1.42e-03 | 4.31e-02 |
| Mitochondrial protein import | 1.43e-03 | 4.31e-02 |
| tRNA processing in the nucleus | 1.46e-03 | 4.31e-02 |
| Pyruvate metabolism and Citric Acid (TCA) cycle | 1.48e-03 | 4.31e-02 |
| Glycosphingolipid metabolism | 1.50e-03 | 4.31e-02 |
| Complex I biogenesis | 1.50e-03 | 4.31e-02 |
| tRNA Aminoacylation | 1.50e-03 | 4.31e-02 |
| tRNA modification in the nucleus and cytosol | 1.52e-03 | 4.31e-02 |
| Voltage gated Potassium channels | 1.52e-03 | 4.31e-02 |
| RNA Polymerase III Transcription Initiation | 1.55e-03 | 4.31e-02 |
| Uptake and actions of bacterial toxins | 1.57e-03 | 4.31e-02 |
| Cristae formation | 1.62e-03 | 4.31e-02 |
| Cytosolic tRNA aminoacylation | 1.63e-03 | 4.31e-02 |
| Cytosolic iron-sulfur cluster assembly | 1.68e-03 | 4.31e-02 |
| Mitochondrial iron-sulfur cluster biogenesis | 1.68e-03 | 4.31e-02 |
| Processing of SMDT1 | 1.70e-03 | 4.31e-02 |
| Regulation of pyruvate dehydrogenase (PDH) complex | 1.70e-03 | 4.31e-02 |
| CDC6 association with the ORC:origin complex | 1.74e-03 | 4.31e-02 |
| Neuronal System | 2.21e-03 | 4.31e-02 |
| Asparagine N-linked glycosylation | 2.28e-03 | 4.31e-02 |
| Metallothioneins bind metals | 2.40e-03 | 4.31e-02 |
| Transport to the Golgi and subsequent modification | 2.55e-03 | 4.31e-02 |
| A tetrasaccharide linker sequence is required for GAG synthesis | 2.58e-03 | 4.31e-02 |
| RUNX2 regulates osteoblast differentiation | 2.58e-03 | 4.31e-02 |
| Sphingolipid metabolism | 2.74e-03 | 4.31e-02 |
| RUNX2 regulates bone development | 2.75e-03 | 4.31e-02 |
| COPI-mediated anterograde transport | 2.75e-03 | 4.31e-02 |
| Signaling by NOTCH3 | 2.94e-03 | 4.31e-02 |
| Packaging Of Telomere Ends | 2.94e-03 | 4.31e-02 |
| Chemokine receptors bind chemokines | 2.94e-03 | 4.31e-02 |
| Interleukin-3, Interleukin-5 and GM-CSF signaling | 2.94e-03 | 4.31e-02 |
| Binding and Uptake of Ligands by Scavenger Receptors | 2.97e-03 | 4.31e-02 |
| Chondroitin sulfate/dermatan sulfate metabolism | 2.99e-03 | 4.31e-02 |

| | | |
|---|---|---|
| Recognition and association of DNA glycosylase with site containing an affected pyrimidine | 3.05e-03 | 4.31e-02 |
| Cleavage of the damaged pyrimidine | 3.05e-03 | 4.31e-02 |
| Depyrimidination | 3.05e-03 | 4.31e-02 |
| Recognition and association of DNA glycosylase with site containing an affected purine | 3.06e-03 | 4.31e-02 |
| Cleavage of the damaged purine | 3.06e-03 | 4.31e-02 |
| Depurination | 3.06e-03 | 4.31e-02 |
| Formation of the ternary complex, and subsequently, the 43S complex | 3.06e-03 | 4.31e-02 |
| RNA Polymerase I Promoter Opening | 3.13e-03 | 4.31e-02 |
| Base-Excision Repair, AP Site Formation | 3.13e-03 | 4.31e-02 |
| Translation initiation complex formation | 3.13e-03 | 4.31e-02 |
| Ribosomal scanning and start codon recognition | 3.13e-03 | 4.31e-02 |
| DNA methylation | 3.14e-03 | 4.31e-02 |
| Activation of the mRNA upon binding of the cap-binding complex and eIFs, and subsequent binding to 43S | 3.14e-03 | 4.31e-02 |
| Activated PKN1 stimulates transcription of AR (androgen receptor) regulated genes KLK2 and KLK3 | 3.25e-03 | 4.31e-02 |
| SIRT1 negatively regulates rRNA expression | 3.27e-03 | 4.31e-02 |
| Citric acid cycle (TCA cycle) | 3.29e-03 | 4.31e-02 |
| Condensation of Prophase Chromosomes | 3.31e-03 | 4.31e-02 |
| DNA Damage/Telomere Stress Induced Senescence | 3.37e-03 | 4.31e-02 |
| Nonhomologous End-Joining (NHEJ) | 3.37e-03 | 4.31e-02 |
| Meiotic recombination | 3.37e-03 | 4.31e-02 |
| RMTs methylate histone arginines | 3.37e-03 | 4.31e-02 |
| Telomere Maintenance | 3.37e-03 | 4.31e-02 |
| ERCC6 (CSB) and EHMT2 (G9a) positively regulate rRNA expression | 3.37e-03 | 4.31e-02 |
| Recruitment and ATM-mediated phosphorylation of repair and signaling proteins at DNA double strand breaks | 3.38e-03 | 4.31e-02 |
| PRC2 methylates histones and DNA | 3.38e-03 | 4.31e-02 |
| Antigen processing: Ubiquitination & Proteasome degradation | 3.41e-03 | 4.31e-02 |
| Deposition of new CENPA-containing nucleosomes at the centromere | 3.42e-03 | 4.31e-02 |
| Nucleosome assembly | 3.42e-03 | 4.31e-02 |
| Meiotic synapsis | 3.44e-03 | 4.31e-02 |
| DNA Double Strand Break Response | 3.45e-03 | 4.31e-02 |
| Formation of the beta-catenin:TCF transactivating complex | 3.46e-03 | 4.31e-02 |
| B-WICH complex positively regulates rRNA expression | 3.46e-03 | 4.31e-02 |
| rRNA processing in the mitochondrion | 3.48e-03 | 4.31e-02 |
| RNA Polymerase I Promoter Escape | 3.50e-03 | 4.31e-02 |
| Base Excision Repair | 3.53e-03 | 4.31e-02 |
| Peptide chain elongation | 3.53e-03 | 4.31e-02 |
| Viral mRNA Translation | 3.53e-03 | 4.31e-02 |
| RUNX1 regulates genes involved in megakaryocyte differentiation and platelet function | 3.57e-03 | 4.31e-02 |
| Neddylation | 3.58e-03 | 4.31e-02 |
| Nonsense Mediated Decay (NMD) independent of the Exon Junction Complex (EJC) | 3.61e-03 | 4.31e-02 |
| Eukaryotic Translation Elongation | 3.62e-03 | 4.31e-02 |
| Selenocysteine synthesis | 3.62e-03 | 4.31e-02 |
| Eukaryotic Translation Termination | 3.62e-03 | 4.31e-02 |
| Pre-NOTCH Transcription and Translation | 3.66e-03 | 4.31e-02 |
| Chromosome Maintenance | 3.68e-03 | 4.31e-02 |
| Negative epigenetic regulation of rRNA expression | 3.68e-03 | 4.31e-02 |
| Positive epigenetic regulation of rRNA expression | 3.68e-03 | 4.31e-02 |
| Interferon gamma signaling | 3.69e-03 | 4.31e-02 |
| RHO GTPases activate PKNs | 3.69e-03 | 4.31e-02 |
| HDACs deacetylate histones | 3.69e-03 | 4.31e-02 |
| GTP hydrolysis and joining of the 60S ribosomal subunit | 3.69e-03 | 4.31e-02 |
| SRP-dependent cotranslational protein targeting to membrane | 3.69e-03 | 4.31e-02 |
| Mitotic G2-G2/M phases | 3.71e-03 | 4.31e-02 |
| Formation of a pool of free 40S subunits | 3.73e-03 | 4.31e-02 |
| Amyloid fiber formation | 3.75e-03 | 4.31e-02 |
| L13a-mediated translational silencing of Ceruloplasmin expression | 3.75e-03 | 4.31e-02 |
| Meiosis | 3.76e-03 | 4.31e-02 |
| Pre-NOTCH Expression and Processing | 3.79e-03 | 4.31e-02 |
| Senescence-Associated Secretory Phenotype (SASP) | 3.79e-03 | 4.31e-02 |
| RNA Polymerase I Promoter Clearance | 3.79e-03 | 4.31e-02 |
| Nonsense-Mediated Decay (NMD) | 3.79e-03 | 4.31e-02 |
| Nonsense Mediated Decay (NMD) enhanced by the Exon Junction Complex (EJC) | 3.79e-03 | 4.31e-02 |

| | | |
|---|---|---|
| RNA Polymerase I Transcription | 3.82e-03 | 4.31e-02 |
| Interleukin-4 and Interleukin-13 signaling | 3.82e-03 | 4.31e-02 |
| NoRC negatively regulates rRNA expression | 3.83e-03 | 4.31e-02 |
| Eukaryotic Translation Initiation | 3.83e-03 | 4.31e-02 |
| Cap-dependent Translation Initiation | 3.83e-03 | 4.31e-02 |
| Immunoregulatory interactions between a Lymphoid and a non-Lymphoid cell | 3.86e-03 | 4.31e-02 |
| Selenoamino acid metabolism | 3.86e-03 | 4.31e-02 |
| S Phase | 3.87e-03 | 4.31e-02 |
| Oxidative Stress Induced Senescence | 3.88e-03 | 4.31e-02 |
| Activation of anterior HOX genes in hindbrain development during early embryogenesis | 3.88e-03 | 4.31e-02 |
| Activation of HOX genes during differentiation | 3.88e-03 | 4.31e-02 |
| RUNX1 regulates transcription of genes involved in differentiation of HSCs | 3.95e-03 | 4.33e-02 |
| Cell surface interactions at the vascular wall | 3.95e-03 | 4.33e-02 |
| Influenza Viral RNA Transcription and Replication | 4.05e-03 | 4.40e-02 |
| Metabolism of nucleotides | 4.13e-03 | 4.45e-02 |
| Influenza Infection | 4.18e-03 | 4.47e-02 |
| HATs acetylate histones | 4.24e-03 | 4.48e-02 |
| Influenza Life Cycle | 4.26e-03 | 4.48e-02 |
| Regulation of expression of SLITs and ROBOs | 4.31e-03 | 4.49e-02 |
| Estrogen-dependent gene expression | 4.33e-03 | 4.49e-02 |
| Major pathway of rRNA processing in the nucleolus and cytosol | 4.59e-03 | 4.72e-02 |
| Peptide ligand-binding receptors | 4.67e-03 | 4.78e-02 |
| PECAM1 interactions | 4.91e-03 | 4.91e-02 |
| G2/M Transition | 4.96e-03 | 4.91e-02 |
| Metabolism of cofactors | 4.98e-03 | 4.91e-02 |
| Defective B4GALT7 causes EDS, progeroid type | 5.01e-03 | 4.91e-02 |
| Defective B3GALT6 causes EDSP2 and SEMDJL1 | 5.01e-03 | 4.91e-02 |
| Regulation of FZD by ubiquitination | 5.01e-03 | 4.91e-02 |
| rRNA processing in the nucleus and cytosol | 5.10e-03 | 4.96e-02 |
| Metabolism of porphyrins | 5.14e-03 | 4.96e-02 |

**Supplementary Table 6 Pathway enrichment of genes ranked by their weight for PLS *model-2* and *component-2*.**

| **Pathway** | **P-value** | **BH-corrected *P*-value** |
|---|---|---|
| Neutrophil degranulation | 1.40e-03 | 1.67e-02 |
| Metabolism of amino acids and derivatives | 1.44e-03 | 1.67e-02 |
| Asparagine N-linked glycosylation | 1.45e-03 | 1.67e-02 |
| Translation | 1.45e-03 | 1.67e-02 |
| Infectious disease | 1.45e-03 | 1.67e-02 |
| HIV Infection | 1.51e-03 | 1.67e-02 |
| Transport to the Golgi and subsequent modification | 1.52e-03 | 1.67e-02 |
| Separation of Sister Chromatids | 1.52e-03 | 1.67e-02 |
| Fatty acid metabolism | 1.52e-03 | 1.67e-02 |
| Neddylation | 1.52e-03 | 1.67e-02 |
| G2/M Transition | 1.53e-03 | 1.67e-02 |
| Mitotic Anaphase | 1.53e-03 | 1.67e-02 |
| Mitotic Metaphase and Anaphase | 1.53e-03 | 1.67e-02 |
| Metabolism of vitamins and cofactors | 1.53e-03 | 1.67e-02 |
| Mitotic G2-G2/M phases | 1.53e-03 | 1.67e-02 |
| Regulation of expression of SLITs and ROBOs | 1.54e-03 | 1.67e-02 |
| The citric acid (TCA) cycle and respiratory electron transport | 1.54e-03 | 1.67e-02 |
| Intra-Golgi and retrograde Golgi-to-ER traffic | 1.54e-03 | 1.67e-02 |
| ER to Golgi Anterograde Transport | 1.55e-03 | 1.67e-02 |
| Protein localization | 1.55e-03 | 1.67e-02 |
| Mitotic G1-G1/S phases | 1.55e-03 | 1.67e-02 |
| C-type lectin receptors (CLRs) | 1.56e-03 | 1.67e-02 |
| PTEN Regulation | 1.56e-03 | 1.67e-02 |
| G1/S Transition | 1.56e-03 | 1.67e-02 |
| S Phase | 1.56e-03 | 1.67e-02 |

| | | |
|---|---|---|
| Beta-catenin independent WNT signaling | 1.56e-03 | 1.67e-02 |
| DNA Replication | 1.56e-03 | 1.67e-02 |
| Signaling by Hedgehog | 1.56e-03 | 1.67e-02 |
| Metabolism of nucleotides | 1.57e-03 | 1.67e-02 |
| COPI-mediated anterograde transport | 1.57e-03 | 1.67e-02 |
| Fc epsilon receptor (FCERI) signaling | 1.57e-03 | 1.67e-02 |
| Antigen processing-Cross presentation | 1.57e-03 | 1.67e-02 |
| Downstream TCR signaling | 1.57e-03 | 1.67e-02 |
| Protein folding | 1.57e-03 | 1.67e-02 |
| TCR signaling | 1.58e-03 | 1.67e-02 |
| CLEC7A (Dectin-1) signaling | 1.58e-03 | 1.67e-02 |
| UCH proteinases | 1.58e-03 | 1.67e-02 |
| COPI-dependent Golgi-to-ER retrograde traffic | 1.58e-03 | 1.67e-02 |
| Golgi-to-ER retrograde transport | 1.58e-03 | 1.67e-02 |
| Mitochondrial translation | 1.58e-03 | 1.67e-02 |
| Host Interactions of HIV factors | 1.58e-03 | 1.67e-02 |
| Respiratory electron transport, ATP synthesis by chemiosmotic coupling, and heat production by uncoupling proteins. | 1.58e-03 | 1.67e-02 |
| Synthesis of DNA | 1.58e-03 | 1.67e-02 |
| TNFR2 non-canonical NF-kB pathway | 1.58e-03 | 1.67e-02 |
| Cyclin A:Cdk2-associated events at S phase entry | 1.59e-03 | 1.67e-02 |
| Hedgehog 'on' state | 1.59e-03 | 1.67e-02 |
| DNA Replication Pre-Initiation | 1.59e-03 | 1.67e-02 |
| MHC class II antigen presentation | 1.60e-03 | 1.67e-02 |
| Metabolism of polyamines | 1.60e-03 | 1.67e-02 |
| Mitochondrial translation elongation | 1.61e-03 | 1.67e-02 |
| APC/C-mediated degradation of cell cycle proteins | 1.61e-03 | 1.67e-02 |
| Regulation of mitotic cell cycle | 1.61e-03 | 1.67e-02 |
| Switching of origins to a post-replicative state | 1.61e-03 | 1.67e-02 |
| Chaperonin-mediated protein folding | 1.61e-03 | 1.67e-02 |
| Mitochondrial translation initiation | 1.61e-03 | 1.67e-02 |
| Mitochondrial translation termination | 1.61e-03 | 1.67e-02 |
| Cyclin E associated events during G1/S transition | 1.61e-03 | 1.67e-02 |
| Downstream signaling events of B Cell Receptor (BCR) | 1.61e-03 | 1.67e-02 |
| ER-Phagosome pathway | 1.61e-03 | 1.67e-02 |
| Regulation of mRNA stability by proteins that bind AU-rich elements | 1.61e-03 | 1.67e-02 |
| Signaling by the B Cell Receptor (BCR) | 1.61e-03 | 1.67e-02 |
| Hedgehog 'off' state | 1.62e-03 | 1.67e-02 |
| Macroautophagy | 1.62e-03 | 1.67e-02 |
| Respiratory electron transport | 1.62e-03 | 1.67e-02 |
| Regulation of APC/C activators between G1/S and early anaphase | 1.62e-03 | 1.67e-02 |
| PCP/CE pathway | 1.62e-03 | 1.67e-02 |
| Regulation of PTEN stability and activity | 1.63e-03 | 1.67e-02 |
| Degradation of beta-catenin by the destruction complex | 1.63e-03 | 1.67e-02 |
| Signaling by NOTCH4 | 1.63e-03 | 1.67e-02 |
| Orc1 removal from chromatin | 1.63e-03 | 1.67e-02 |
| Mitochondrial protein import | 1.64e-03 | 1.67e-02 |
| Oxygen-dependent proline hydroxylation of Hypoxia-inducible Factor Alpha | 1.64e-03 | 1.67e-02 |
| Regulation of RAS by GAPs | 1.64e-03 | 1.67e-02 |
| G1/S DNA Damage Checkpoints | 1.64e-03 | 1.67e-02 |
| Activation of NF-kappaB in B cells | 1.64e-03 | 1.67e-02 |
| APC/C:Cdc20 mediated degradation of Securin | 1.64e-03 | 1.67e-02 |
| Assembly of the pre-replicative complex | 1.64e-03 | 1.67e-02 |
| ABC transporter disorders | 1.64e-03 | 1.67e-02 |
| FCERI mediated NF-kB activation | 1.64e-03 | 1.67e-02 |
| Activation of APC/C and APC/C:Cdc20 mediated degradation of mitotic proteins | 1.64e-03 | 1.67e-02 |
| rRNA modification in the nucleus and cytosol | 1.65e-03 | 1.67e-02 |
| p53-Dependent G1 DNA Damage Response | 1.65e-03 | 1.67e-02 |
| p53-Dependent G1/S DNA damage checkpoint | 1.65e-03 | 1.67e-02 |
| APC:Cdc20 mediated degradation of cell cycle proteins prior to satisfation of the cell cycle checkpoint | 1.65e-03 | 1.67e-02 |
| Hedgehog ligand biogenesis | 1.65e-03 | 1.67e-02 |
| Defective CFTR causes cystic fibrosis | 1.65e-03 | 1.67e-02 |
| Regulation of RUNX2 expression and activity | 1.65e-03 | 1.67e-02 |
| CDK-mediated phosphorylation and removal of Cdc6 | 1.66e-03 | 1.67e-02 |

| | | |
|---|---|---|
| APC/C:Cdh1 mediated degradation of Cdc20 and other APC/C:Cdh1 targeted proteins in late mitosis/early G1 | 1.66e-03 | 1.67e-02 |
| Cdc20:Phospho-APC/C mediated degradation of Cyclin A | 1.66e-03 | 1.67e-02 |
| Cellular response to hypoxia | 1.66e-03 | 1.67e-02 |
| APC/C:Cdc20 mediated degradation of mitotic proteins | 1.66e-03 | 1.67e-02 |
| The role of GTSE1 in G2/M progression after G2 checkpoint | 1.66e-03 | 1.67e-02 |
| SCF(Skp2)-mediated degradation of p27/p21 | 1.66e-03 | 1.67e-02 |
| Autodegradation of Cdh1 by Cdh1:APC/C | 1.66e-03 | 1.67e-02 |
| Dectin-1 mediated noncanonical NF-kB signaling | 1.66e-03 | 1.67e-02 |
| Degradation of GLI1 by the proteasome | 1.66e-03 | 1.67e-02 |
| Degradation of GLI2 by the proteasome | 1.66e-03 | 1.67e-02 |
| GLI3 is processed to GLI3R by the proteasome | 1.66e-03 | 1.67e-02 |
| Asymmetric localization of PCP proteins | 1.67e-03 | 1.67e-02 |
| AUF1 (hnRNP D0) binds and destabilizes mRNA | 1.67e-03 | 1.67e-02 |
| Hh mutants that don't undergo autocatalytic processing are degraded by ERAD | 1.67e-03 | 1.67e-02 |
| CDT1 association with the CDC6:ORC:origin complex | 1.67e-03 | 1.67e-02 |
| Stabilization of p53 | 1.67e-03 | 1.67e-02 |
| Iron uptake and transport | 1.67e-03 | 1.67e-02 |
| Pyruvate metabolism and Citric Acid (TCA) cycle | 1.67e-03 | 1.67e-02 |
| Hh mutants abrogate ligand secretion | 1.68e-03 | 1.67e-02 |
| NIK-->noncanonical NF-kB signaling | 1.68e-03 | 1.67e-02 |
| Vif-mediated degradation of APOBEC3G | 1.68e-03 | 1.67e-02 |
| FBXL7 down-regulates AURKA during mitotic entry and in early mitosis | 1.68e-03 | 1.67e-02 |
| Negative regulation of NOTCH4 signaling | 1.68e-03 | 1.67e-02 |
| HSP90 chaperone cycle for steroid hormone receptors (SHR) | 1.69e-03 | 1.67e-02 |
| Regulation of Apoptosis | 1.69e-03 | 1.67e-02 |
| SCF-beta-TrCP mediated degradation of Emi1 | 1.69e-03 | 1.67e-02 |
| Degradation of AXIN | 1.69e-03 | 1.67e-02 |
| Regulation of RUNX3 expression and activity | 1.69e-03 | 1.67e-02 |
| Autodegradation of the E3 ubiquitin ligase COP1 | 1.69e-03 | 1.67e-02 |
| Complex I biogenesis | 1.69e-03 | 1.67e-02 |
| Cross-presentation of soluble exogenous antigens (endosomes) | 1.69e-03 | 1.67e-02 |
| Regulation of activated PAK-2p34 by proteasome mediated degradation | 1.69e-03 | 1.67e-02 |
| Degradation of DVL | 1.69e-03 | 1.67e-02 |
| Recycling pathway of L1 | 1.70e-03 | 1.67e-02 |
| Ubiquitin-dependent degradation of Cyclin D | 1.71e-03 | 1.67e-02 |
| Vpu mediated degradation of CD4 | 1.71e-03 | 1.67e-02 |
| Regulation of ornithine decarboxylase (ODC) | 1.71e-03 | 1.67e-02 |
| Ubiquitin Mediated Degradation of Phosphorylated Cdc25A | 1.71e-03 | 1.67e-02 |
| p53-Independent DNA Damage Response | 1.71e-03 | 1.67e-02 |
| p53-Independent G1/S DNA damage checkpoint | 1.71e-03 | 1.67e-02 |
| Gap junction trafficking and regulation | 1.71e-03 | 1.67e-02 |
| Gluconeogenesis | 1.72e-03 | 1.67e-02 |
| ROS and RNS production in phagocytes | 1.72e-03 | 1.67e-02 |
| COPI-independent Golgi-to-ER retrograde traffic | 1.72e-03 | 1.67e-02 |
| Gap junction trafficking | 1.73e-03 | 1.67e-02 |
| DNA Damage Recognition in GG-NER | 1.74e-03 | 1.67e-02 |
| Gap junction assembly | 1.74e-03 | 1.67e-02 |
| Carboxyterminal post-translational modifications of tubulin | 1.76e-03 | 1.67e-02 |
| Signaling by FGFR2 IIIa TM | 1.76e-03 | 1.67e-02 |
| RNA Pol II CTD phosphorylation and interaction with CE during HIV infection | 1.77e-03 | 1.67e-02 |
| RNA Pol II CTD phosphorylation and interaction with CE | 1.77e-03 | 1.67e-02 |
| Activation of AMPK downstream of NMDARs | 1.77e-03 | 1.67e-02 |
| Transferrin endocytosis and recycling | 1.77e-03 | 1.67e-02 |
| Cooperation of Prefoldin and TriC/CCT in actin and tubulin folding | 1.77e-03 | 1.67e-02 |
| RHO GTPases activate IQGAPs | 1.77e-03 | 1.67e-02 |
| Cristae formation | 1.77e-03 | 1.67e-02 |
| mRNA Capping | 1.78e-03 | 1.67e-02 |
| Prefoldin mediated transfer of substrate to CCT/TriC | 1.78e-03 | 1.67e-02 |
| Post-chaperonin tubulin folding pathway | 1.80e-03 | 1.67e-02 |
| Abortive elongation of HIV-1 transcript in the absence of Tat | 1.81e-03 | 1.67e-02 |
| Insulin receptor recycling | 1.81e-03 | 1.67e-02 |
| Nucleotide salvage | 1.83e-03 | 1.67e-02 |
| Formation of tubulin folding intermediates by CCT/TriC | 1.83e-03 | 1.67e-02 |

| | | |
|---|---|---|
| Citric acid cycle (TCA cycle) | 1.83e-03 | 1.67e-02 |
| Microtubule-dependent trafficking of connexons from Golgi to the plasma membrane | 1.86e-03 | 1.69e-02 |
| Transport of connexons to the plasma membrane | 1.88e-03 | 1.69e-02 |
| Pregnenolone biosynthesis | 1.91e-03 | 1.71e-02 |
| mitochondrial fatty acid beta-oxidation of saturated fatty acids | 1.96e-03 | 1.74e-02 |
| Formation of Senescence-Associated Heterochromatin Foci (SAHF) | 2.13e-03 | 1.89e-02 |
| Regulation of MECP2 expression and activity | 2.35e-03 | 2.07e-02 |
| HDMs demethylate histones | 2.43e-03 | 2.12e-02 |
| Activated PKN1 stimulates transcription of AR (androgen receptor) regulated genes KLK2 and KLK3 | 2.49e-03 | 2.13e-02 |
| Transcriptional Regulation by MECP2 | 2.49e-03 | 2.13e-02 |
| SIRT1 negatively regulates rRNA expression | 2.52e-03 | 2.13e-02 |
| DNA methylation | 2.52e-03 | 2.13e-02 |
| Meiotic synapsis | 2.54e-03 | 2.13e-02 |
| PKMTs methylate histone lysines | 2.54e-03 | 2.13e-02 |
| DNA Damage/Telomere Stress Induced Senescence | 2.56e-03 | 2.13e-02 |
| SUMOylation of chromatin organization proteins | 2.57e-03 | 2.13e-02 |
| PRC2 methylates histones and DNA | 2.57e-03 | 2.13e-02 |
| Pre-NOTCH Expression and Processing | 2.58e-03 | 2.13e-02 |
| Formation of the beta-catenin:TCF transactivating complex | 2.62e-03 | 2.14e-02 |
| Pre-NOTCH Transcription and Translation | 2.64e-03 | 2.14e-02 |
| Gene Silencing by RNA | 2.73e-03 | 2.20e-02 |
| HATs acetylate histones | 2.74e-03 | 2.20e-02 |
| Mitotic Prophase | 2.79e-03 | 2.22e-02 |
| Estrogen-dependent gene expression | 2.80e-03 | 2.22e-02 |
| SUMOylation | 2.86e-03 | 2.25e-02 |
| Cellular Senescence | 2.87e-03 | 2.25e-02 |
| SUMO E3 ligases SUMOylate target proteins | 2.90e-03 | 2.26e-02 |
| Rab regulation of trafficking | 3.16e-03 | 2.43e-02 |
| L1CAM interactions | 3.18e-03 | 2.43e-02 |
| Resolution of Sister Chromatid Cohesion | 3.19e-03 | 2.43e-02 |
| Chromatin modifying enzymes | 3.21e-03 | 2.43e-02 |
| Chromatin organization | 3.21e-03 | 2.43e-02 |
| MAPK6/MAPK4 signaling | 3.22e-03 | 2.43e-02 |
| mRNA Splicing - Minor Pathway | 3.37e-03 | 2.53e-02 |
| Metabolism of steroid hormones | 3.43e-03 | 2.56e-02 |
| Formation of the Early Elongation Complex | 3.52e-03 | 2.60e-02 |
| Formation of the HIV-1 Early Elongation Complex | 3.52e-03 | 2.60e-02 |
| Cholesterol biosynthesis | 3.55e-03 | 2.61e-02 |
| Diseases associated with glycosylation precursor biosynthesis | 3.59e-03 | 2.63e-02 |
| Cytosolic tRNA aminoacylation | 3.62e-03 | 2.64e-02 |
| Metabolism of cofactors | 3.70e-03 | 2.68e-02 |
| Formation of ATP by chemiosmotic coupling | 3.75e-03 | 2.70e-02 |
| CDC6 association with the ORC:origin complex | 3.81e-03 | 2.73e-02 |
| Tight junction interactions | 4.54e-03 | 3.23e-02 |
| NOTCH1 Intracellular Domain Regulates Transcription | 4.78e-03 | 3.39e-02 |
| Metabolism of water-soluble vitamins and cofactors | 4.82e-03 | 3.40e-02 |
| trans-Golgi Network Vesicle Budding | 4.97e-03 | 3.48e-02 |
| RNA Polymerase I Promoter Opening | 4.99e-03 | 3.48e-02 |
| ERCC6 (CSB) and EHMT2 (G9a) positively regulate rRNA expression | 5.04e-03 | 3.50e-02 |
| RMTs methylate histone arginines | 5.13e-03 | 3.55e-02 |
| Mitochondrial Fatty Acid Beta-Oxidation | 5.24e-03 | 3.60e-02 |
| Assembly and cell surface presentation of NMDA receptors | 5.26e-03 | 3.60e-02 |
| Assembly of active LPL and LIPC lipase complexes | 5.39e-03 | 3.66e-02 |
| Oxidative Stress Induced Senescence | 5.39e-03 | 3.66e-02 |
| RUNX1 regulates genes involved in megakaryocyte differentiation and platelet function | 5.42e-03 | 3.66e-02 |
| Epigenetic regulation of gene expression | 5.48e-03 | 3.68e-02 |
| Phenylalanine and tyrosine catabolism | 5.87e-03 | 3.92e-02 |
| Interleukin-1 signaling | 6.28e-03 | 4.17e-02 |
| Visual phototransduction | 6.32e-03 | 4.17e-02 |
| Mitochondrial biogenesis | 6.33e-03 | 4.17e-02 |
| Apoptosis induced DNA fragmentation | 6.44e-03 | 4.20e-02 |
| Regulation of IFNG signaling | 6.44e-03 | 4.20e-02 |
| COPII-mediated vesicle transport | 6.53e-03 | 4.24e-02 |
| Translocation of SLC2A4 (GLUT4) to the plasma membrane | 6.58e-03 | 4.25e-02 |

| | | |
|---|---|---|
| Golgi Associated Vesicle Biogenesis | 6.78e-03 | 4.36e-02 |
| Notch-HLH transcription pathway | 6.82e-03 | 4.36e-02 |
| tRNA Aminoacylation | 6.84e-03 | 4.36e-02 |
| Detoxification of Reactive Oxygen Species | 6.94e-03 | 4.41e-02 |
| Packaging Of Telomere Ends | 7.28e-03 | 4.60e-02 |
| Platelet activation, signaling and aggregation | 7.32e-03 | 4.60e-02 |
| Metabolism of porphyrins | 7.43e-03 | 4.66e-02 |
| Mitotic Prometaphase | 7.65e-03 | 4.76e-02 |
| SUMOylation of DNA damage response and repair proteins | 7.69e-03 | 4.76e-02 |
| Deposition of new CENPA-containing nucleosomes at the centromere | 7.75e-03 | 4.76e-02 |
| Nucleosome assembly | 7.75e-03 | 4.76e-02 |
| RHO GTPases Activate Formins | 7.79e-03 | 4.76e-02 |
| Interleukin-1 family signaling | 7.81e-03 | 4.76e-02 |
| Meiosis | 8.09e-03 | 4.89e-02 |
| RAB GEFs exchange GTP for GDP on RABs | 8.09e-03 | 4.89e-02 |
